# Synergy between vesicular and non-vesicular gliotransmission regulates synaptic plasticity and working memory

**DOI:** 10.1101/2021.03.25.437028

**Authors:** Ulyana Lalo, Seyed Rasooli-Nejad, Alexander Bogdanov, Lorenzo More, Wuhyun Koh, Jurgen Muller, Mark Wall, C. Justin Lee, Yuriy Pankratov

## Abstract

Astrocytes are an active element of brain signalling, capable of release of small molecule gliotransmitters by vesicular and channel-mediated mechanisms. However, specific physiological roles of astroglial exocytosis of glutamate and D-Serine remain controversial. Our data demonstrate that cortical astrocytes can release glutamate and D-Serine by combination of SNARE-dependent exocytosis and non-vesicular mechanisms dependent on TREK-1 and Best1 channels. Astrocyte-derived glutamate and D-serine elicited complex multicomponent phasic response in neocortical pyramidal neurons, which is mediated by extra-synaptic GluN2B receptors. Impairment of either pathway of gliotransmission (in the TREK1 KO, Best-1 KO or dnSNARE mice) strongly affected the NMDAR-dependent long-term synaptic plasticity in the hippocampus and neocortex. Moreover, impairment of astroglial exocytosis in dnSNARE mice led to the deficit in the spatial working memory which was rescued by environmental enrichment.

We conclude that synergism between vesicular and non-vesicular gliotransmission is crucial for astrocyte-neuron communication and astroglia-driven regulation of synaptic plasticity and memory.

**Highlights:**

- Astrocytes *in situ* release glutamate via exocytosis and channel-mediated release.
- Astroglia-derived glutamate and D-Serine activate phasic NMDAR currents in neurons
- Both vesicular and non-vesicular gliotransmission are required for synaptic plasticity
- Impaired exocytosis of gliotransmitters causes deficit in working memory

## INTRODUCTION

The perception of the role of glial cells in brain function has undergone a revolutionary change in recent years and the research field of glia-neuron interaction is rapidly expanding. There is a growing evidence that glia, in particular astrocytes, can receive and integrate information on neural activity and can respond by increasing the metabolic support for neurons and modulating synaptic transmission via release of gliotransmitters (Araque et al., 2014; Attwell et al., 2010; Gourine et al., 2010; Halassa and Haydon, 2010; Lalo et al., 2014a). As in the many developing research areas, many controversies and unresolved questions have arisen particularly in how glia and neurons communicate (Araque et al., 2014; Bazargani and Attwell, 2016; Savtchouk and Volterra, 2018). Arguably, one of the most debated issues is the mechanism and physiological relevance of Ca^2+^-dependent release of gliotransmitters from astrocytes (Bazargani and Attwell, 2016; Fiacco and McCarthy, 2018; Savtchouk and Volterra, 2018). There are three prevalent views on this topic: 1) astrocytes can release gliotransmitters, in particular ATP, Glutamate and D-Serine, via Ca^2+^- and SNARE-dependent exocytosis (Bezzi et al., 2004; Covelo and Araque, 2018; Henneberger et al., 2010; Lalo et al., 2014b; Pascual et al., 2005; Rasooli-Nejad et al., 2014; Schwarz et al., 2017; Sultan et al., 2015); 2) gliotransmitters can be released via Ca^2+^-dependent (Fields and Burnstock, 2006; Woo et al., 2012); 3) Ca^2+^- and SNARE-dependent gliotransmission does not play an important role in regulation of synaptic function (Agulhon et al., 2008; Fiacco and McCarthy, 2018; Fujita et al., 2014). In addition, the possibility that gliotransmitters can be released of via Ca^2+^-independent channels, has also been reported (Woo et al., 2012, Num et al. 2019). Thus, it still remains elusive what mechanism, vesicular or channel-mediated, plays predominant role in the release of glutamatergic transmitters from astrocytes and glial modulation of cognitive functions.

It should also be noted that the majority of experimental results on the release of glutamatergic gliotransmitters have been obtained from cultured astrocytes and data directly showing neuronal responses to astrocyte-derived transmitters *in situ* are scarce (Lalo et al., 2014a; Woo et al., 2012; Nam et al.2019). The data on the cognitive effects of vesicular release of gliotransmitters are also scarce. Despite growing evidence that glial modulation of synaptic plasticity is impaired in brain slice from dn-SNARE mice (Lalo et al., 2014a; Pankratov and Lalo, 2015; Pascual et al., 2005), the behavioral memory-related phenotype in these mice has been yet to be reported. Furthermore, the importance of astrocytes as a source of D-Serine – the crucial NMDA receptor co-agonist – has recently ignited a strong debate (Papouin et al., 2017b; Wolosker et al., 2016). Thus it is clear that the topic of glutamatergic glia-neuron interactions still remains highly contentious.

It has been rightfully noted recently (Araque et al., 2014; Bazargani and Attwell, 2016; Savtchouk and Volterra, 2018) that now it is time for next wave of research into glia physiology – research aimed to resolve the controversies and to unify the conflicting views. Our previous results (Lalo et al., 2014a; Woo et al., 2012) presented strong evidence to support both sides of the “vesicular vs non-vesicular gliotransmission” argument, so here we have endeavored to resolve the conflicting views on the fundamental mechanisms of gliotransmission. We have explored the synergy between exocytotic and non-exocytotic release of gliotransmitters, in particular glutamate and D-Serine. In this paper we present data which, we believe, can bridge the great divide in the field. Our electrophysiological and behavioural data demonstrate that vesicular and non-vesicular gliotransmission work together in the regulation of synaptic plasticity and memory.

## RESULTS

### Vesicular and non-vesicular release of glutamate from acutely isolated cortical astrocytes

As a first step in the analysis of the mechanism of Ca^2+^-dependent release of glutamate from astrocytes we used a sniffer-cell approach where glutamate was detected by HEK293 cells transfected with mutant GluR1-L497Y receptors (Woo et al., 2012). These receptors have much higher affinity to glutamate (EC_50_ around 10 µM) and desensitize much slower than native GluR1 receptors and thus are suitable as glutamate-detectors. Cortical astrocytes were dissociated with the aid of a non-enzymatic isolation procedure (see *Methods*) and thus were devoid of the cell culturing artifacts (Lalo et al., 2014a; Lalo et al., 2006). We compared the release of glutamate from astrocytes of somatosensory cortex of wild-type mice and from mice astroglia-specific inducible expression of dominant-negative SNARE (dnSNARE mice).

Living dissociated cells were immunolabeled with antibodies to specific astroglial markers and vesicular glutamate transporters ((Lalo et al., 2014a); see also Methods) and then distributed into a recording chamber containing pre-plated HEK293-GluR1-L497Y cells (Fig.1). In some experiments, astrocytes were also pre-loaded with Ca^2+^-indicator Calcium Green-2AM and glial cell marker SR-101. To evaluate the contribution of Ca^2+^-dependent vesicular and non-vesicular mechanisms, isolated astrocytes were separately pre-incubated with BAPTA-AM or with the inhibitor of vesicular glutamate transporters (VGLUTs) Rose Bengal (see *Methods*).

Whole-cell voltage-clamp recording were made from a HEK293-GluR1-L497Y cells (Fig.1A,B) laying in very close proximity to astrocytes, identified initially by immunolabeling (wild-type mice) or by EGFP fluorescence (in dn-SNARE mice) and later by their electrophysiological properties (Lalo et al., 2014a; Lalo et al., 2006). We elevated the cytosolic Ca^2+^ concentration in the astrocytes by applying agonists of PAR-1 receptors, CB1 endocannabinoid receptors or α1-adrenoreceptors (Fig. 1B, FigS1). The reliability and specificity of this method to activate astrocytes has been confirmed in our previous experiments (Lalo et al., 2014a; Pankratov and Lalo, 2015; Rasooli-Nejad et al., 2014; Woo et al., 2012, Oh et al., 2019).

**Figure 1.**
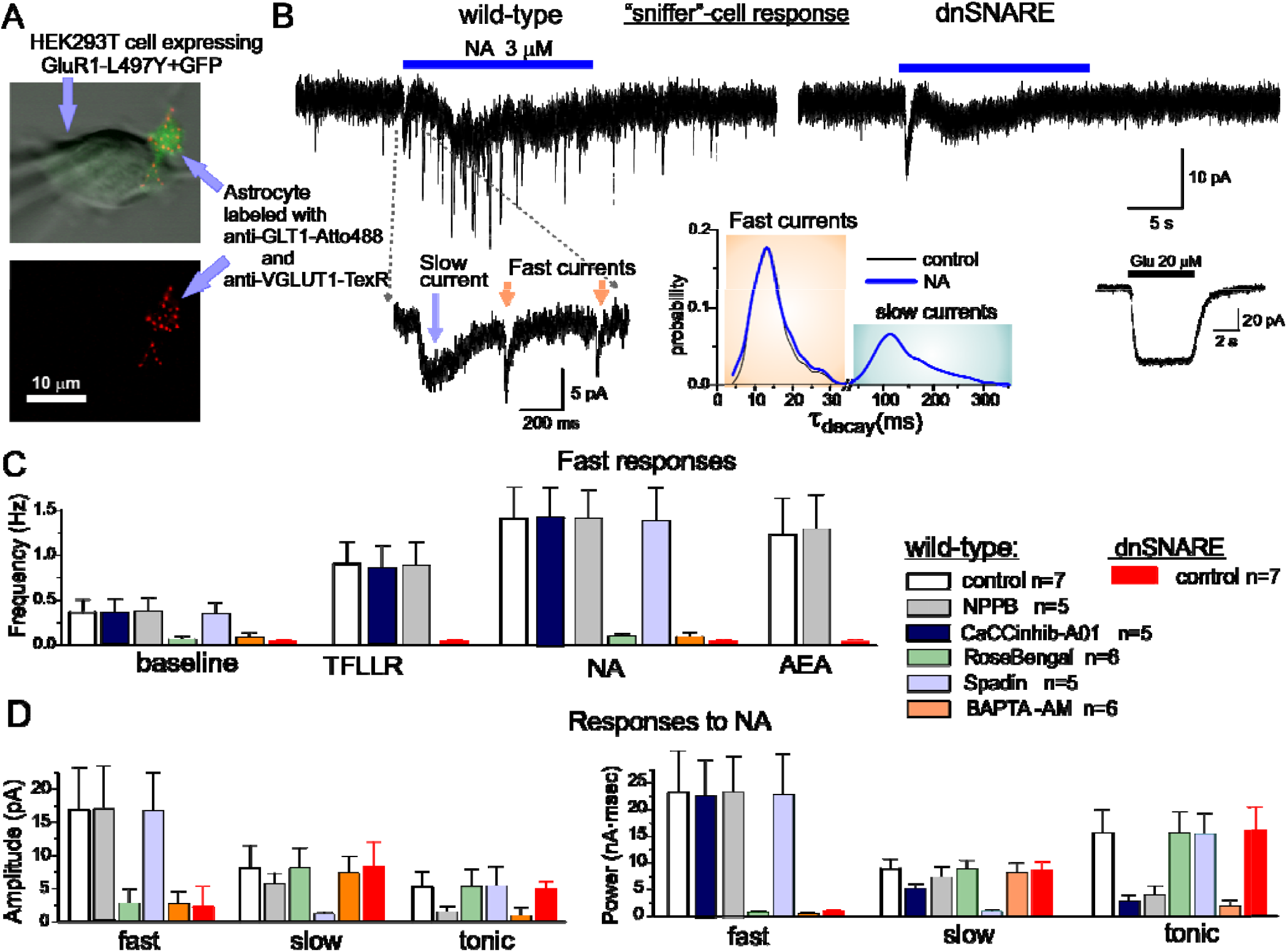
Glutamatergic gliotransmission evaluated by “sniffer”-cell approach. **(A)** Acutely dissociated cortical astrocytes were placed on top HEK293 cells expressing GluR1- L497Y glutamate receptors. Punctate staining of astrocytes with VGLUT1 antibodies suggests the presence of glutamate-containing vesicles. **(B)** Whole-cell recording of transmembrane current in the HEK293-GluR1-L497Y cells at -80 mV. Noradrenaline-mediated Ca^2+^-elevation in the astrocytes induced three kinds of responses in the HEK293-GluR1-L497Y: fast transient “synaptic-like” currents and slower transient currents, both shown in the inset, and slow tonic current. Decay time constant distribution of transient glia-induced currents shows two distinct groups of events. The fast currents were not elicited by dnSNARE-expressing astrocytes suggesting their vesicular origin. *Inset*, response of sniffer cell to 20 μM glutamate **(C)** Average frequency of fast glutamatergic responses activated via different astrocytic receptors under various conditions: wild-type astrocytes in control, after pre-incubation of astrocytes with BAPTA-AM or VGLUT inhibitor Rose Bengal (500 nM) and in the presence of Best1 channel blockers NPPB (30 µM) and CaCCinhib-A01 (20 µM), and TREK-1 selective inhibitor spadin (200 nM); and dnSNARE astro ytes in control. **(D)** Pooled data on the average amplitude and overall charge transferred by the three types of glutamatergic currents elicited after activation of wild-type and dnSNARE astrocytes via NA-receptors. The tonic current was dramatically reduced by NPPB and CaCCinhib-A01, the slow transient current was unaffected by the dnSNARE expression but was eliminated by spadin, blocker of TREK-1 channels. Fast transient currents were significantly inhibited by the VGLUT inhibitor, BAPTA-AM and astroglial expression of dnSNARE. Combined, these data suggest that cortical astrocytes can release glutamate by vesicular and non-vesicular mechanisms.

Activation of astrocytes by 30 sec-long applications of noradrenaline (NA, 3 µM), PAR-1 agonist TFLLR (10 µM) or CB1 agonist anandamide (AEA, 250 µM) induced complex responses in the adjacent detector cells (Fig1B,C). Three components of the responses could be clearly distinguished based on their kinetics and functional properties: (1) fast transient currents with amplitudes of 5-20 pA and decay time constants of 7-25 ms which resembled the parameters of exocytosis-activated synaptic currents (Canal et al., 2011; Espinosa and Kavalali, 2009; McAllister and Stevens, 2000; Pankratov et al., 2007) and currents evoked by exocytosis of ATP from astrocytes (Lalo et al., 2014a); (2) slow transient currents with decay time of 100-250 ms and (3) tonic changes in the baseline whole-cell current which lasted for 1-5 s (Fig. 1B). Similar to previous reports (Woo et al. 2012), the HEK293-GluR1-L497Y cells, which did not have adjacent astrocytes, did not show any response to either NA (n=8) or TFLLR (n=5). This allows us to attribute the multi-component response solely to the astrocytic release of glutamate. Importantly, astrocytes from dnSNARE mice did not produce the fast responses in the detector cells in all 21 experiments. These results suggest that the fast glia-induced pulsatile currents have vesicular origin. In contrast, astrocytes from dnSNARE mice produced the slow transient and tonic components of response in all 21 experiments. Activation of slow and tonic responses by dnSNARE astrocytes strongly suggested the lack of side-effects of dnSNARE expression on the trafficking of astroglial membrane proteins, which was also supported by our previous data (Lalo et al., 2014a; Rasooli-Nejad et al., 2014) and further substantiated by the data on the lack of effect of dnSNARE expression in the activity of Best1 channels in cortical astrocytes (Supplementary Figure S1).

All astrocytes tested were able to induce the tonic current, with the slow transient current observed in approximately in 68% of trials (8 out 13 WT and 8 out of 12 dn-SNARE astrocytes). The burst of fast pulsatile glutamatergic current was observed in 80% of experiments with WT astrocytes. Similarly to currents induced by vesicular release of ATP (Lalo et al., 2014a), pulsatile fast glutamatergic currents could be also observed in baseline conditions but with much lower frequency. In contrast, the slow transient and tonic currents appeared only after activation of astrocytes. The pharmacological properties of the different components of astrocyte-induced sniffer-cell responses corroborated the notion of different molecular mechanisms of their generation. Firstly, both the frequency and amplitude of the fast transient currents were markedly (up to 72 and 74%) reduced by the 20 min long pre-incubation of wild-type astrocytes with the inhibitor of vesicular glutamate transporters Rose Bengal (Fig.1C,D). At the same time, neither the slow transient nor tonic responses were affected by the VGLUT inhibitor. The fast currents were also dramatically inhibited by pre-incubation of the astrocytes with Ca^2+^-chelator BAPTA-AM (Fig.1C,D). Based on their functional properties, the fast sniffer-cell currents can be attributed to the Ca^2+^-dependent exocytosis of glutamate from astrocytes with a high degree of confidence.

In contrast to the fast currents, the slow transient responses were not affected by BAPTA but were significantly inhibited by spadin (200 nM), a selective antagonist of TREK-1 potassium channels (Borsotto et al., 2015), and only moderately affected by NPPB (30 µM), an inhibitor of Ca^2+^-activated chloride channels (CAAC). These data suggest that the slow transient responses originate from non-vesicular Ca^2+^-independent release of glutamate via TREK-1 channels. Involvement of this pathway in glutamate release has been previously reported for hippocampal astrocytes (Woo et al., 2012, Nam et al, 2019). The tonic component of the response was substantially reduced after pre-incubation of astrocytes with BAPTA (Fig.1C,D), indicating its Ca^2+^-dependent nature. The tonic currents were not sensitive to spadin but were strongly inhibited by NPPB. It has been reported that NPPB has rather weak selectivity for CAAC and also inhibits some two-pore potassium channels. However at the concentration used (30 µM) it should preferentially inhibit CAACs (Greenwood and Leblanc, 2007). The novel, more specific CAAC inhibitor (De La Fuente et al., 2008) CACCInh-A01 (20 µM) also inhibited the tonic component by 71±11%, n=5, (Fig.1C,D). The kinetics and pharmacological sensitivity of the slow transient and tonic currents induced by activation of neocortical astrocytes closely agrees with our previous observations (Woo et al., 2012) that release of glutamate from hippocampal astrocytes occurs via TREK-1-containing K2P potassium channels and Best1 CAACs denoted correspondingly as “fast” and “slow” currents (Woo et al., 2012).

Taken together, our data demonstrate that neocortical astrocytes are capable of the activity-dependent release of glutamate via three alternative mechanism: Ca^2+^- and SNARE-dependent vesicular release, Ca^2+^-dependent non-vesicular release via chloride channels and Ca^2+^-independent release via TREK-1 potassium channels. The relative contribution for each mechanism in NA-activated release of glutamate, evaluated as net charge transferred by glutamatergic currents in detector cells (Fig.1D), was 48%, 33% and 19% correspondingly for exocytosis (fast transient currents), non-vesicular Ca^2+^-dependent (tonic currents) and non-vesicular Ca^2+^ independent (slow transient currents). Thus, the above results suggest that both vesicular and non-vesicular mechanisms can be important for glutamatergic gliotransmission.

### Release of glutamate from astrocytes *in situ* occurs via SNARE-dependent and non-vesicular pathways

The putative physiological effects of glutamatergic gliotransmitters are supposedly mediated by NMDA receptors abundantly expressed by pyramidal neurons in many brain areas (Paoletti et al., 2013; Papouin and Oliet, 2014; Sanz-Clemente et al., 2013), both at synaptic densities and extrasynaptic sites. To assess the neuronal response to glia-derived glutamate we recorded the whole-cell NMDAR-mediated currents in neocortical pyramidal neurons in presence of TTX at membrane potential of -40 mV and physiological Mg^2+^concentration (Fig.2). Activation of AMPA, GABA and P2X receptors was inhibited by DNQX (30 µM), picrotoxin (100 µM) and PPADS (10 µM) plus 5-BDBD (5 µM). The selective activation of astrocytes (Lalo et al., 2014a; Pankratov and Lalo, 2015; Woo et al., 2012) was achieved with a 30 sec-long application of either noradrenaline (3 µM) or the PAR-1 agonist TFLLR (10 µM).

As suggested by the “sniffer”-cell experiments, one could expect the multi-modal glial release of glutamate via vesicular and non-vesicular pathways which might activate neuronal responses of different kinetics and functional properties. In order to inhibit astroglial exocytosis of glutamate, we perfused individual astrocytes with an intracellular solution containing 3 nM tetanus neurotoxin light chain (TeNTx) and 30 μM of the calcium indicator Calcium Green-2 (Fig.2A) and recorded mEPSCs in a neighbouring neuron (lying within 30 µm distance from the perfused astrocyte). In the second line of experiments, we perfused astrocytes with an intracellular solution containing 3 mM BAPTA (instead of TeNTX) to clamp down Ca^2+^ elevations. Perfusion of astrocytes with a solution only containing Calcium Green-2 was used as a control. The electrophysiological recordings in neurons started 10-15 min after perfusion of astrocytes.

Under baseline conditions (before the activation of astrocytes), the miniature spontaneous synaptic currents (mEPSCs) were observed in all neurons tested (Fig.2B). The amplitude of baseline spontaneous inward currents in wild-type mice was 5.4±1.9 pA (n=25) and the decay time constant was 25.4±8.9 ms (Fig 2C, S2). In each individual neuron, the baseline spontaneous currents exhibited wide and skewed distribution of the decay time with main peak at 20-25 ms accompanied by significant fraction of events with decay time about 30-50 ms (Fig 2C, S2).

**Figure 2.**
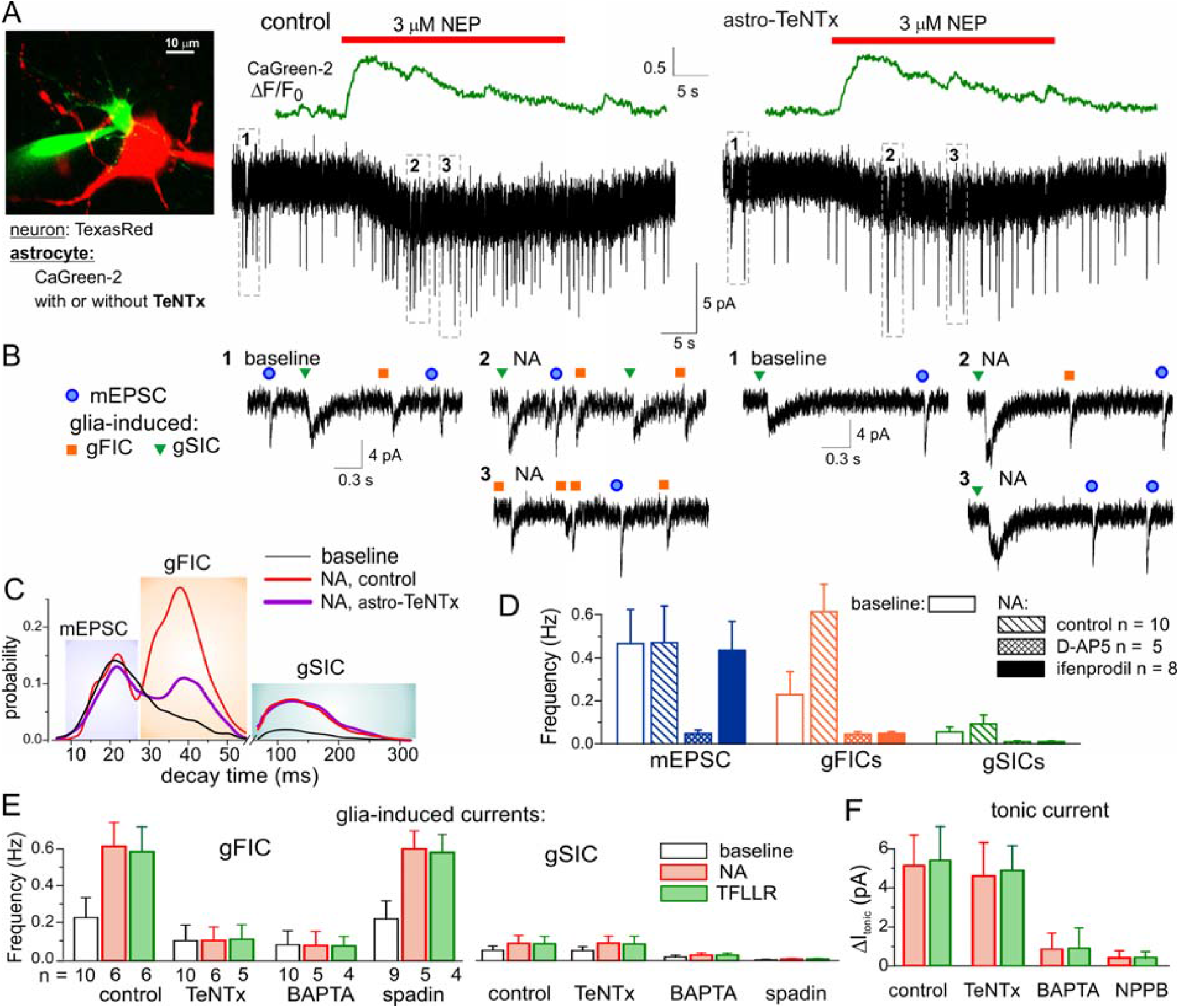
Release of glutamate from astrocytes in the neocortical slices. (A) Whole-cell currents were recorded in the pyramidal neuron of layer 2/3 of neocortical slice at -40 mV in the presence of picrotoxin, TTX, DNQX, PPADS and 5-BDBD simultaneously with perfusion of an astrocyte with intracellular solution containing either fluorescent dye Ca-Green-2 alone or Ca-Green-2 and 3nM of Tetanus neurotoxin light chain or 3 mM BAPTA. Activation of astrocytic Ca^2+^signalling via PAR1 or α1-adrenoreceptors elicited the burst of transient NMDAR-mediated currents and caused a slow inward deflection in the holding current (tonic response) in the neurons. (B-D) The transient currents can be subdivided into three groups by their functional properties. (B) examples of individual currents recorded in period indicated on graph in the panel (A). (C) the decay time distributions of transient currents recorded before (baseline) and after stimulation of astrocytes with noradrenaline. *Left peak*: The frequency of currents with decay time around 20 ms was not strongly affected by stimulation of astrocytes; these currents were denoted as mEPSCs. *Middle peak*: the frequency of currents with decay time around 40-45 ms significantly increased after stimulation of astrocytes but this increase was attenuated by perfusion of astrocyte with TeNTX; these currents were denoted as glia-induced faster currents (gFICs). *Right peak*: the frequency of transient currents with decay time around 100-200 ms significantly increased after stimulation of astrocytes but was not affected by inhibition of astrocytic exocytosis with TeNTX; these currents were denoted as glia-induced slower currents (gSICs) (D) Pooled data (mean±SD) on the frequency of three types of transient currents before (baseline) and after stimulation of astrocytes with noradrenaline in the control and in the presence of GluN2B inhibitor ifenprodil(3 μM). Note that only gFICs and gSICs were affected by ifenprodil. (E) Pooled data on the changes in the frequency of gFICs and gSICs induced after activation of astrocytes agonists of PAR1 (TFLLR) or α1-adrenoreceptors (NA) in control and after perfusion of individual astrocytes with TeNTX or BAPTA or inhibition of TREK-1 channels by spadin. The frequency of gFICs was strongly reduced by TeNTX but not spadin whereas gSICs showed the opposite pattern. (F) The pooled data on the amplitude of tonic current induced after activation of astrocytes with noradrenaline and TFLLR. The glia-induced tonic current was not affected by TeNTX but was inhibited by BAPTA and inhibitor of Best1 channels NPPB.

Activation of astrocytic Ca^2+^-transients led to the appearance of a large number of transient inward currents which could be distinguished from the main fraction of baseline mEPSCs by their slower kinetics (Fig.2C). The transient glia-induced currents were also accompanied by a very slow inward deflection of the holding current (Fig.2B) referred further as “tonic response”.

The decay time distribution of transient currents recorded after activation of astrocytes exhibited three clear components: 1) fast currents with decay time constant of 22.7±5.5 ms (left peak) which coincided with parameters of the baseline mEPSCs; 2) fast currents with decay time of 43.8±9.7 ms (middle peak), this fraction was significantly reduced by perfusion of astrocytes with TeNTX; and 3) slow currents with decay time constant of 155±63 ms (right peak). Neither activation of astrocytes nor perfusion with TeNTX altered the frequency and statistical weight of currents of the group 1 verifying their synaptic origin (Fig.2C,D). In contrast, the frequency of currents in groups 2 and 3 increased dramatically after activation of astrocytes, their decay kinetics were close to those transient currents recorded in the “sniffer”-cells. This strongly suggested their origin from gliotransmission. For clarity, we denoted the currents of group 2 and 3 as glia-induced fast and slow inward currents; correspondingly gFICs and gSICs.

It is very likely that glutamate released from astrocytes at extrasynaptic cites would preferentially activate extrasynaptic NMDA receptors. The extrasynaptic NMDARs expressed in many brain areas, including neocortex, contain the GluN2B subunit in contrast to synaptic receptors (Paoletti et al., 2013; Papouin and Oliet, 2014) which predominantly contain GluN2A subunits. Thus, the neuronal currents activated by synaptic and the glial release of glutamate could have different sensitivities to the specific GluN2B antagonist ifenprodil. Thus we tested the pharmacological sensitivity of the three groups of NA-induced currents. Indeed, the fast currents identified preliminary as mEPSCs, were not sensitive to ifenprodil, in contrast to the gFICs and gSICs (Fig.2D) which frequencies was decreased correspondingly by 93±7% and 90±5% (n=8). All components of response to NA were eliminated by D-AP5 and thus all currents were mediated by NMDARs.

In addition to the difference in the kinetics, the average amplitude of gFICs was smaller than the quantal amplitude of mEPSCs (Fig.S2). The different biophysical and pharmacological properties of mEPSCs and gFICs strongly support their different origin. The astroglial origin of gFICs was corroborated by the data obtained in the dnSNARE mice (Fig.S2) where the frequency of ifenprodil-sensitive currents of slower kinetics and smaller amplitude was much less compared to the wild-type mice.

To verify the origin of gFICs and gSICs we evaluated the effect of intracellular perfusion of astrocytes with TeNTX and BAPTA and the effect of spadin, an inhibitor of TREK1 channels (Fig.2E). The frequency of gFICs recorded in the neurons located in the vicinity of astrocytes perfused with TeNTx or BAPTA (Fig.2A) was significantly reduced as compared to control, both in the baseline conditions and after application of NA or TFLLR (Fig.2E). At the same time, the gFICs were not sensitive to spadin. The combination of the functional and pharmacological properties of gFICs (Fig.2, Fig.S2) allows us to unequivocally attribute them to the SNARE-dependent exocytosis of glutamate from astrocytes.

In contrast to gFICs, the glia-induced slow inward currents (gSICs) were not affected by intracellular perfusion of astrocytes with TeNTx but were eliminated by application of spadin (Fig.2E) strongly supporting their origin from astroglial TREK1-mediated release of glutamate. The amplitude of tonic response to activation of astrocytes was significantly reduced by perfusion of astrocytes with BAPTA but was only marginally decreased by TeNTx or spadin (Fig.2B,F); tonic response was also efficiently inhibited by NPPB (Fig.2F) and ifenprodil (by 93.4±5.9%, n=8, data not shown). These result support the main contribution of CACC–mediated release to glia-induced tonic current.

Our data on functional and pharmacological properties of the three components of glia-induced response in neurons *in situ* (Fig.2) are in a good agreement with properties of astrocyte-induced glutamatergic currents in the “sniffer”–cells (Fig.1). Combined, these data strongly support the capability of astrocytes to simultaneously release glutamate via vesicular and non-vesicular mechanisms. To elucidate further the putative roles for the different non-vesicular mechanisms of glial release of glutamate, we evaluated the glia-induced inwards currents in the neurons of TREK-1 and Best1 knockout mice and compared them to currents recorded in the wild-type mice (Fig.3). Genetic deletion of TREK-1 channels did not affect the fast glia-induced currents but almost completely eliminated the slow currents (Fig. 3A,B,D). On average, the overall frequency of gSICs was 95±6% (n=6) lower in the TREK-1 KO mice as compared to the wild-type mice. In 50% of trials, we observed only a few gSICs in neurons from TREK-1 KO mice over all experimental period (40-45 min); the other 50% of neurons did not show any gSICs at all. This result provides a compelling evidence for a predominant role for TREK-1 channel-mediated release in the generation of gSICs. The lack of changes in the gFICs in the TREK-1 KO mice closely agrees with our data (Fig.2, S2) on the origin of gFICs from the glial exocytosis of glutamate. We also noted some changes in the gSICs in the Best1 KO mice (Fig.1A3) which were related to the deficit of D-Serine as described below.

**Figure 3.**
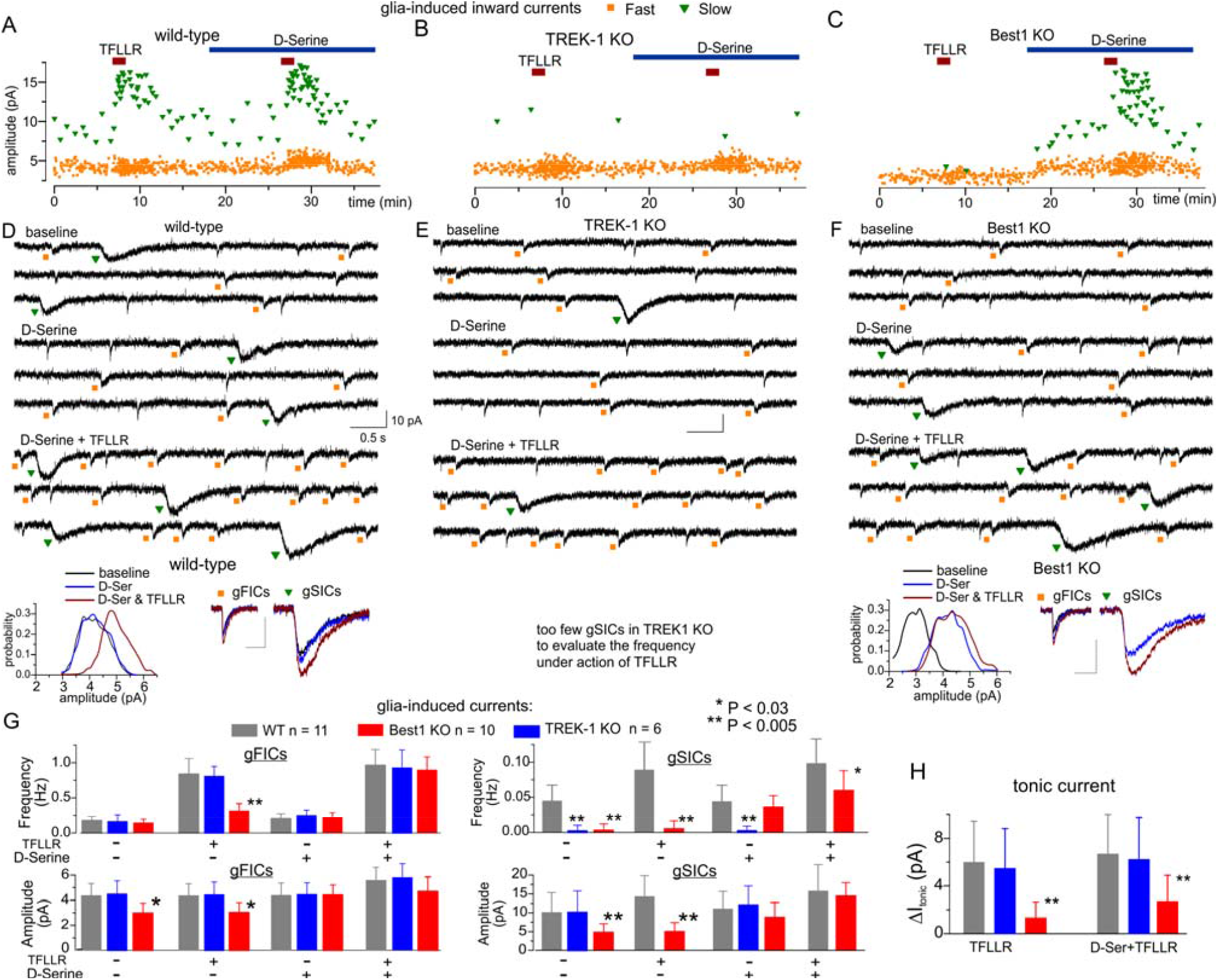
Role for Best1 channels in glutamatergic gliotransmission in the neocortex. The transient glia-induced currents were recorded in neocortical pyramidal neurons of wild-type and Best1-KO mice using the same experimental paradigm as in Fig.2. Astrocyte Ca^2+^ -signalling was selectively activated by the PAR-1 receptor agonist TFLLR. (A-C) the time course of glia-induced fast (gFICs) and slow (gSICs) currents recorded in the neocortical pyramidal neurons of wild-type (A), TREK-1 (B) and Best1 KO (C) mice. The fast and slow glia-induced currents were discriminated by their kinetics as shown in Fig.2. (D-F) The corresponding examples of neuronal currents, average waveforms and amplitude histograms of gFICs and gSICs recorded in control, after application of exogenous D-serine and activation of astrocytes by TFLLR in presence of D-Serine. (G,H) The pooled data (mean±SD) on the frequency and amplitude of glia-induced fast (gFICs) and slow (gSICs) currents and the amplitude of the tonic TFLLR-induced current measured as in Fig.2A,F. The asterisks (* and **) indicate the statistical significance of the difference between WT and transgenic mice. In the wild-type mice, activation of astrocytes with TFLLR markedly increased the frequency of gFICs but both the amplitude and frequency of gSICs. The exogenous D-Serine does not have significant effect on glia-induced currents in WT mice. Note almost complete cessation of gSICs in the TREK-1 KO mice both under baseline conditions and in the presence of TFLLR and/or D-Serine. In the Best1 KO mice, the baseline frequency and amplitude of gSICs are significantly reduced but are restored by exogenous D-Serine. The tonic current is unaffected in the TREK-1 KO mice but is significantly reduced in the Best1KO both in the absence and presence of D-Serine. There is no decrease in the frequency of gFICs in the both transgenic mice and only moderate decrease in their amplitude in Best1 KO, the amplitude of gFICs was restored by exogenous D-Serine. These data verify the origin of gSICs and tonic current from glial release of glutamate correspondingly via TREK-1 and Best1 channels. The rescue of gFICs and gSICs phenotype in the Best1KO mice by exogenous D-Serine strongly support the importance of Ca-activated chloride channels for release of D-Serine from astrocytes.

### Best1 Ca^2+^-activated channels bring major contribution to release of D-Serine from cortical astrocytes

In addition to glutamate, astrocytes can also release D-Serine, a crucial co-agonist of NMDA receptors, in a Ca^2+^-dependent manner (Henneberger et al., 2010; Papouin et al., 2017a; Rasooli-Nejad et al., 2014; Sultan et al., 2015). Although vesicular exocytosis is traditionally considered to be the main pathway of glial release of D-Serine (Halassa et al., 2007; Henneberger et al., 2010), participation of non-vesicular mechanisms cannot be excluded *a priori*. As a first evidence of this, we noticed that the slow, spadin-sensitive, glia-induced inward currents in neurons were inhibited after perfusion of astrocytes with BAPTA (Fig.2E) whereas spadin-sensitive slow inward currents in “sniffer”-cells were not affected (Fig.1D). The most parsimonious explanation could be the significant decrease in the concentration of D-Serine which would be critical for NMDAR-mediated gSICs recorded in neurons in slices but not for GluR1-mediated currents in the “sniffer-cell”. This putative decrease in the ambient concentration of D-Serine in the neocortical tissue was caused by perfusion of astrocytes with BAPTA which could affect both Ca^2+^-dependent chloride channels and Ca^2+^-dependent vesicular release but not TeNTx which would affect only exocytosis. This may imply a significant contribution of CACCs in the glial release of D-Serine, which in turn works synergistically with TREK1-mediated release of glutamate in generation of gSICs. This notion was further corroborated by the effects of genetic deletion of Best1 channels on the glia-induced currents (Fig.3C,F,G).

We observed marked changes in the gSICs in the Best1 KO and wild-type mice. First, the average gSICs frequency in the neurons of Best1 KO mice was extremely low in the baseline conditions and showed only moderate increase after activation of glial PAR-1 receptors (dFig.3A,B). The average gSICs amplitude was significantly lower in the Best1 KO as compared to wild-type mice, suggesting that low frequency of gSICs in Best1 KO mice could be caused by falling of most events below the detection threshold. Application of the exogenous D-Serine (10 µM) to the neocortical slices from Best1 KOs mice changed the gSICs dramatically: their amplitude and frequency significantly increased and they become responsive to PAR-1 agonist (Fig.3C). The existence of gSICs in the Best1 KO mice, albeit only in presence of exogenous D-Serine, supports our conclusion that they originate from the release of glutamate via TREK-1 rather than CACC channels. Still, the gSICs critically depend on D-Serine, released via Best1 CACCs.

The other components of glia-evoked response in the neurons of Best1 KO mice were also altered. Although the baseline frequency of fast glia-induced currents did not undergo significant changes, their amplitude was markedly lower than in the wild-type mice. Under control conditions, the TFLLR-induced burst of the gFICs in the Best1 KO mice was less prominent (Fig.3C,F): their frequency increased only by 117±31% as compared to 386±99% increase in the wild-type (Fig.3A,D). Exogenous D-Serine did not affect the baseline frequency of gFICs in the neurons of Best1 KO mice but significantly increased their amplitude, this effect manifested in the right-ward shift in the amplitude histogram (Fig.3F). Exogenous D-Serine restored the effect of TFLLR on the gFICs frequency in the Best1KO mice.

### Astrocytes not neurons play a major role in D-serine release

So, we observed a general trend for all components of glia-evoked responses: there was strong decrease in their power in the Best1 KO mice but phenotype was rescued by the application of exogenous D-Serine. We would like to emphasize that, in contrast to the Best1 KO, exogenous D-Serine did not have any marked effects on all components of glia-evoked responses in the wild-type mice (Fig.3A,G). Combined, the above data strongly support the major contribution of astroglial CACCs into glial release of D-Serine. Our data also show that all three types of GluN2B-mediated neuronal currents (gSICs, gFICs and tonic current) strongly rely on the presence of extracellular D-Serine, this goes in line with recent results (Ferreira et al., 2017).

Our results closely agree with previous data on the major role of astrocytes in the release of D-Serine (Panatier et al., 2006; Papouin et al., 2017b; Sultan et al., 2015). Yet, the importance and specificity of glial release of D-Serine has been questioned recently and a major role for neurons in the release of D-Serine has been speculated instead (Wolosker et al., 2016). Due to the increasing importance of this issue, we directly addressed a putative contribution of neurons in the release of D-serine. To dissect the specific contribution of neurons and astrocytes, we used a preparation of acutely-isolated neocortical neurons which were devoid of the influence of glial cells (Figure S3). We used a technique of non-enzymatic vibro-dissociation which retains functional synapses on the dendrites of isolated neurons, which can be verified by the presence of miniature spontaneous synaptic currents (Duguid et al., 2007; Lalo et al., 2016; Lalo and Pankratov, 2017; Rasooli-Nejad et al., 2014).

We recorded NMDA receptor-mediated currents in the acutely-dissociated neocortical pyramidal neurons at membrane potential of -40 mV in the presence of 100 µM picrotoxin, 30 µM NBQX and 20 µM PPADS. To compensate for the any depletion of neuronal D-Serine content due to intracellular perfusion, intracellular solution was supplemented with 1 mM D-Serine; intracellular concentration of EGTA was set to 0.1 mM. In the absence of external D-Serine or glycine, isolated neurons exhibited a very low baseline frequency of NMDAR-mediated mEPSCs and the application of TFLLR or NA and did not cause notable changes in their amplitude or frequency (Fig. S3A,C). Application of exogenous D-Serine dramatically increased the average amplitude and notably increased the frequency of mEPSCs. In contrast to the isolated neurons, neuronal mEPSCs (insensitive to ifenprodil) recorded in the brain slices, showed only the moderate increase (29±8%, n=7) in the amplitude upon application of exogenous D-Serine (Fig. S3B,C). Activation of astrocytes by TFLLR or NE produced almost the similar increase (25±7%, n=5) in the mEPSCs amplitude which was not enhanced further in the presence of exogenous D-Serine (Fig. S3B,C). Also, our evaluation of D-Serine concentration in the cortical tissues with aid of microelectrode sensors (Fig. S3E) showed that TFLLR-activated transients of D-Serine were not sensitive to inhibition of neuronal activity with CNQX and TTX but were significantly decreased after 15 min-long incubation of slices with glial metabolic poison fluoroacetate (FAC, 3 mM). Combined together, our data strongly support the major role of astrocytes in the activity-dependent release of D-Serine, going in line with recent data (Papouin et al., 2017b; Papouin and Oliet, 2014).

We also would like to note that all kinds of glia-induced neuronal glutamatergic currents showed dependence on Best1 channel-mediated release of D-Serine, although to a different extent. While gSICs almost completely disappeared in the neurons of Best1 KO mice under baseline conditions, the gFICs were only partially inhibited. The most feasible explanation is that generation of gFICs can occur at areas of neuronal membrane exposed to the ambient D-Serine coming from another source. This source could be provided by glial exocytosis of D-Serine, which is supported by our previous data (Rasooli-Nejad et al., 2014) and data of other groups (Papouin et al., 2017a; Sultan et al., 2015). The presence of an ambient extrasynaptic glycine (Papouin et al., 2012) is also possible. Yet, the gFICs strongly rely on release of D-Serine via CACCs as evidenced by their lower amplitude in the Best1 KO mice (Fig.3D). Our results strongly suggest that glycine or D-Serine of vesicular origin can be viewed as merely a supplementary source of NMDAR co-agonists. Dependence of all the components of glutamatergic glia-evoked responses on release of D-Serine via glial CACCs bears important physiological implications, which will be addressed below.

### Synaptically–activated release of glutamate from astrocytes

One of the most intriguing issues regarding a physiological role for gliotransmission is capability of astrocytes to release gliotransmitters in response to the local synaptic activity. We have shown previously that short episodes of high-frequency stimulation of cortical afferents (HFS) are able to activate substantial Ca^2+^-transients in the neocortical astrocytes leading to glial exocytosis of ATP and activation of P2X receptors in pyramidal neurons (Lalo et al., 2014a; Rasooli-Nejad et al., 2014). In the present work, we followed the similar experimental paradigm but instead evaluated the NMDAR-mediated currents in the neocortical neurons (Fig.4).

Under baseline conditions, we observed three types of phasic NMDAR-mediated currents which have been identified as mEPSCs, gFICs and gSICs as described above (Fig.2,3). Short HFS episode activated substantial Ca^2+^ transients in the neocortical astrocytes of both wild-type and dn-SNARE mice (Fig.4A) which were followed by the bursts of phasic NMDAR currents and the appearance of the tonic response (Fig.4B).

The burst of phasic NMDAR activity was much more prominent in the wild-type as opposed to the dn-SNARE mice, suggesting a glial mechanism for this effect. The detailed analysis of NMDAR-currents kinetics showed that it was the gFICs which underwent the HFS-induced enhancement (Fig.4C,E). The increase in the gFICs frequency reached 201±48 % in the wild-type mice. The mEPSCs did not exhibit significant increase in the frequency. The statistical weight of mEPSCs in the overall population of phasic NMDAR-currents did not change (Fig.4C,D). The HFS-induced enhancement of gFICs was impaired in dn-SNARE mice (Fig. 4B,C). Furthermore, the burst of the gFICs activity in the wild type mice was strongly attenuated after perfusion of astrocytes with the VGLUT inhibitor Rose Bengal or BAPTA (Fig.4D,E). These data support our conclusion on their origin from glial exocytosis.

**Figure 4.**
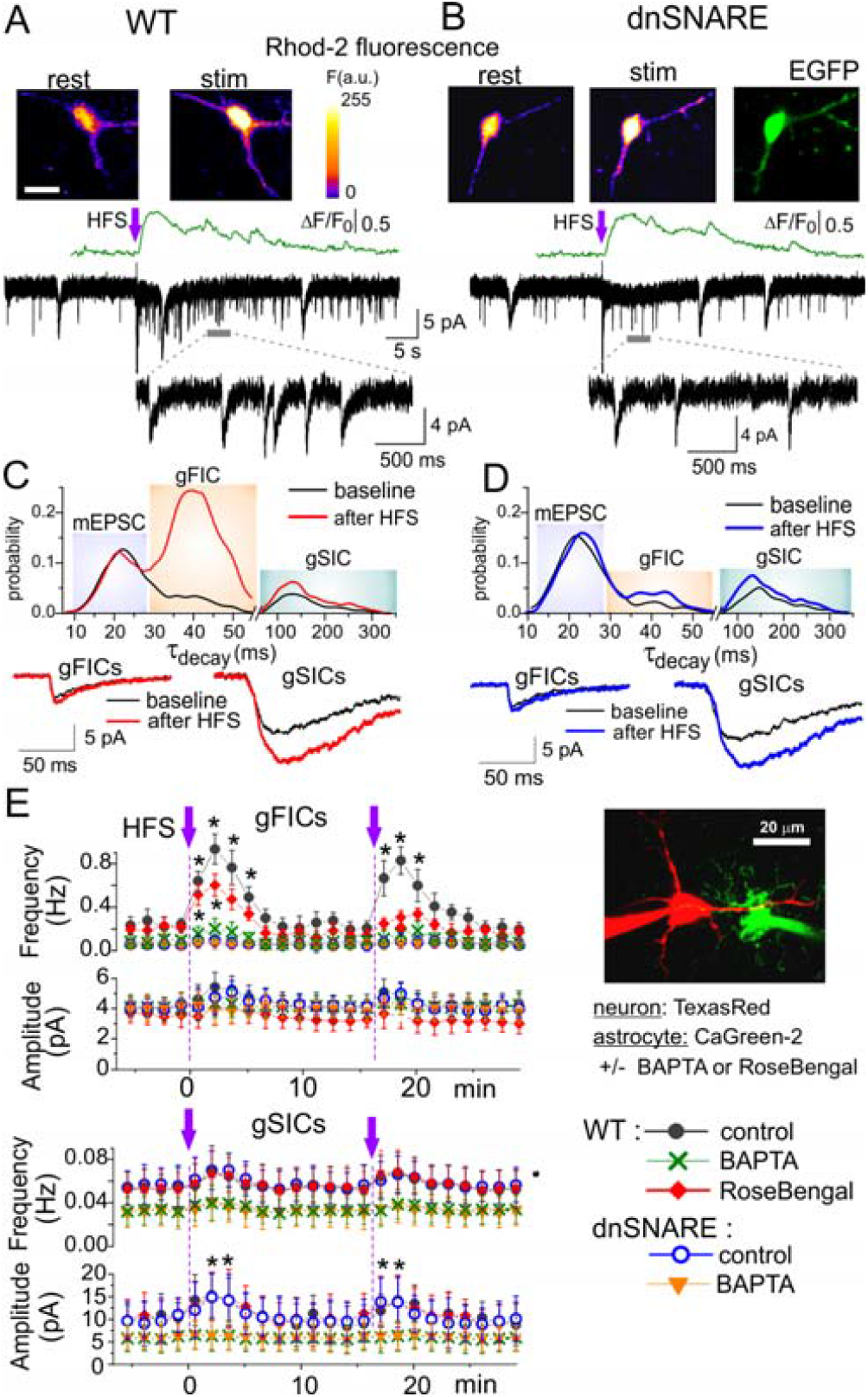
Release of glutamate from astrocytes *in situ* can be triggered by synaptic stimulation. (A,B) Ca^2+^-signalling was monitored in astrocytes (loaded with Rhod-2AM) of somatosensory cortex layer 2/3 of wild-type (A) and dn-SNARE (B) mice simultaneously with voltage-clamp recordings from pyramidal neurons. The NMDAR-mediated spontaneous were recorded in the pyramidal neurons at holding potential of -40 mV in the presence of picrotoxin (100µM), DNQX (50 µM) and P2X antagonists PPADS (10 µM) and 5-BDBD (50 M). The single episode of 100 Hz stimulation (HFS) triggered Ca^2+^-transients in the astrocytes of both wild-type and dn-SNARE mice. (A,B) the Ca^2+^-transients and pseudo-colour images (scale bar 10 μm) recorded before (rest) and at the peak of response (stim). Ca^2+^-transients were followed by burst of transient NMDAR-currents of diverse kinetics, similar to currents activated by application of TFLLR or NEP (Fig.2). (C, D) the decay time distributions of transient currents recorded before (baseline) and after stimulation show the presence of three populations, similar to NEP-activated currents (Fig.2): mEPSCs (decay time ∼20 ms), gFICs (decay time ∼ 40-45 ms) and gSICs (decay time ∼ 100-200 ms). The corresponding waveforms (average of 20 traces) are shown in the inlay. The proportion of gFICs, depicted as middle peak at the distributions, dramatically increased after stimulation in the wild-type (C) but not the dnSNARE mice (D); 5 neurons were tested for each genotype. The proportion of gSICs underwent only moderate increase both in the WT and dnSNARE mice. The frequency of mEPSCs did not show notable changes. (E). The time course of the amplitude and frequency of gFICs (upper graph) and gSICs (lower graph) recorded in the neocortical pyramidal neurons during dialysis of neighbouring astrocytes with intracellular solution containing either fluorescent dye Calcium Green-2 alone (control) or Calcium Green-2 and 100 nM of VGLUT inhibitor Rose Bengal or 3 mM BAPTA. Each dot shows the average value for currents recorded in 1 min time window; data are presented as mean±SD for 7 neurons. The asterisks (*) and (**) indicate the significant difference from the baseline values. The HFS was delivered twice with 15 min interval. Note the significant increase in the gFICs frequency in the control in wild-type but not the dnSNARE mice. The second HFS-evoked increase was reduced by perfusion of neighbouring astrocyte with VGUT inhibitor; perfusion with BAPTA had even stronger effect on gFICs. The amplitude of gSICs showed statistically significant increase both in the WT and dnSNARE mice in the control, this effect was prevented by perfusion of astrocytes with BAPTA.

In contrast to gFICs, the increase in the frequency of gSICs was moderate and reached just 29±12% (Fig.4E). This was accompanied, however, by the significant increase in the amplitude of gSICs both in the wild-type and dn-SNARE mice (Fig.4C-E). The frequency of gSICs was not affected by dn-SNARE expression and perfusion of astrocytes with Rose Bengal but perfusion of astrocytes with BAPTA decreased the amplitude of gSICs and its HFS-induced enhancement (Fig.4E). This behavior agrees with the notion that gSICs are generated by synergetic action of TREK-1 mediated release of glutamate and Ca^2+^-dependent release of D-Serine via Best1 channels (Fig.2,3).The tonic current was observed both in the wild-type and dn-SNARE mice and was sensitive only to perfusion of astrocytes with BAPTA (Fig.4E).

These results closely agree with our data on physiological and pharmacological properties of glia-evoked responses (Fig.2,3) and demonstrate that release of glutamate via vesicular, TREK-1 and Best1-mediated pathways can be activated by physiologically-attainable activation of astrocytes *in situ*. Our data also suggest that exocytosis- and Best-1 channel-dependent mechanisms bring the major contribution to release of glutamate triggered by synaptically-activated astroglial Ca^2+^-signaling.

### Role for vesicular and non-vesicular gliotransmission in the long-term synaptic plasticity

The activity-dependent release of glutamate and D-Serine from astrocytes that, in turn, activates neuronal extrasynaptic NMDA receptors (Figs. 2-4) can underlie the impact of astroglia on synaptic plasticity in the neocortex and hippocampus. In addition to glutamatergic transmitters, astrocytes can also exocytose ATP which, as we have shown previously (Lalo et al., 2014a), can down-regulate tonic and phasic GABAergic inhibition. Since a decrease in the tonic inhibition would contribute towards postsynaptic depolarization and also enhance NMDAR activity (Casasola et al., 2004; Hess et al., 1996; Shen et al., 2010), one would expect the convergence of action purinergic and glutamatergic gliotransmission to enhance the induction of long-term synaptic plasticity. Previous works implicated Ca^2+^-dependent glial release of ATP and D-Serine in the control of synaptic plasticity in the hippocampus (Henneberger et al., 2010; Pascual et al., 2005). The role of astroglia-derived glutamate in the synaptic plasticity has been reported in the hippocampus (Park et al, 2015; Nam et al., 2019). However, its role in the neocortex remains almost unexplored.

We have investigated the long-term potentiation (LTP) of the field excitatory postsynaptic potentials (fEPSP) in the layer II/III of somatosensory cortex of wild-type mice and mice with impaired vesicular and non-vesicular release of gliotransmitters. The fEPSPs were evoked by the stimulation of the same neuronal afferents as EPSCs described above (see Methods). Potentiation of fEPSPs was induced by 5 episodes of theta-burst stimulation (5 TBS). Such stimulation protocol reliably induced LTP in the neocortical slices of wild-type mice; the magnitude of LTP reached 162±12% (n=21) at 60±5 min after induction under control conditions (Figure 5A). The magnitude of LTP was significantly lower when fEPSPs were recorded in the vicinity of astrocytes perfused with TeNTx or an inhibitor of vesicular glutamate transporters (similarly to experiments shown in the Fig2A and 4C). This result demonstrates the importance of astroglial exocytosis of gliotransmitters, in particular glutamate, for LTP induction. One should note that perfusion of astrocytes with TeNTx had a much larger effect than perfusion with a VGLUT inhibitor, suggesting the important contribution of non-glutamatergic gliotransmitters, most likely ATP. Inhibition of putative neuronal targets of glia-derived glutamate and ATP, namely GluN2B (Fig.2-4) and P2X receptors (Lalo et al., 2014a), significantly attenuated the magnitude of LTP (Fig.5B).

**Figure 5.**
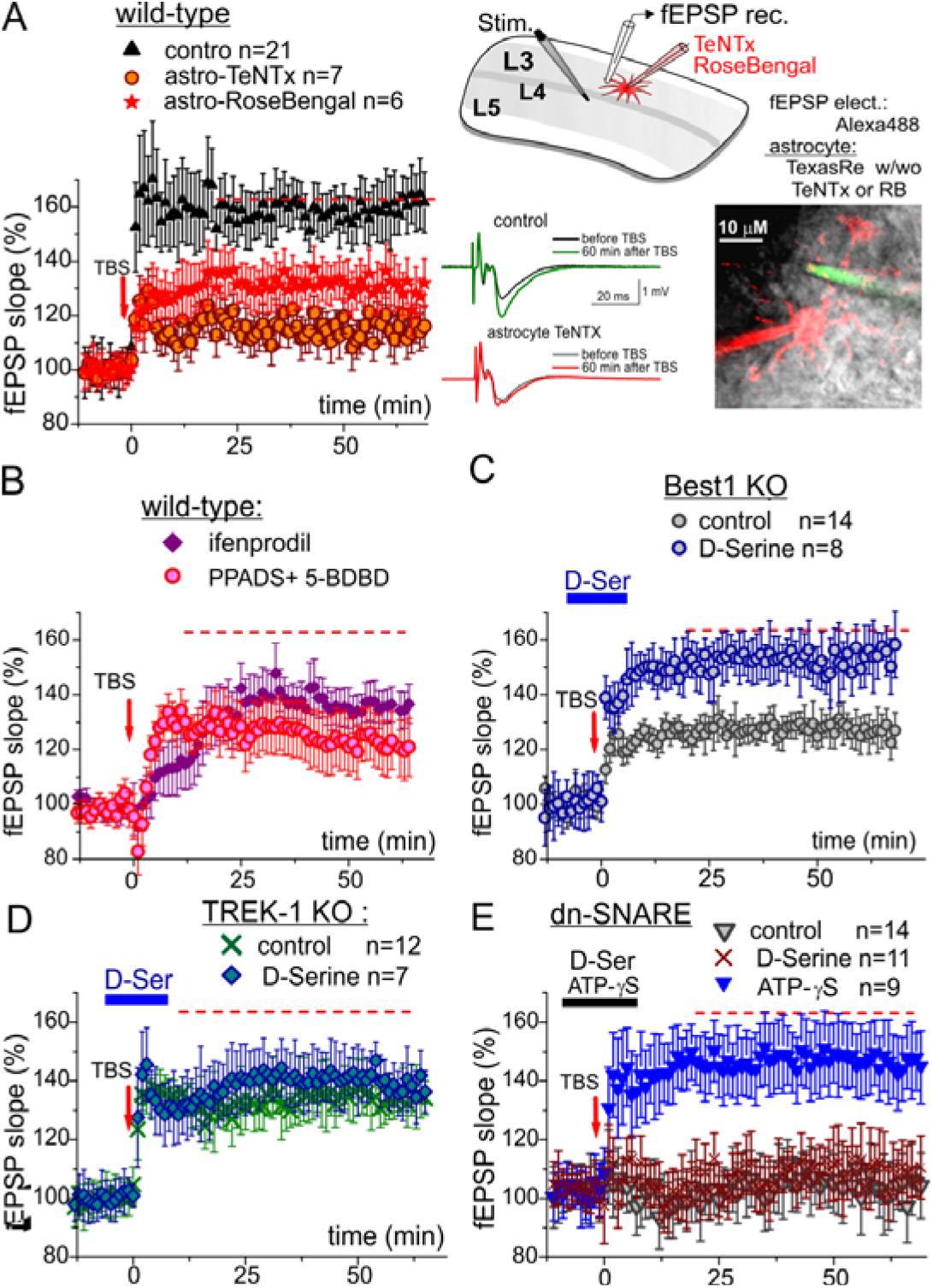
The synergism between different mechanisms of gliotransmission is essential for synaptic plasticity. (A-E) The long-term potentiation of fEPSPs in the neocortical layer 2/3 was induced by 5 theta-burst of high-frequency stimulation in the wild-type, Best1 KO, TREK-1 KO and dnSNARE mice under different conditions: control (the average level for WT mice is indicated as red line in all graphs), intracellular perfusion of individual astrocyte, lying in the vicinity of recording site, with TetanusToxin light chain or VGLUT inhibitor Rose-Bengal (A), antagonists of neuronal NMDAR and P2X receptors (B) and application of exogenous gliotransmitters D-Ser (C-E) and ATP-γS (E) for 5 min prior and 5 min after delivering high-frequency stimulation. (A) Inhibition of the vesicular release of gliotransmitters from individual astrocytes significantly reduced the magnitude of LTP. (B) Inhibition of neuronal GluN2B and P2X receptors, which can be activated by glutamate and ATP released from astrocytes, significantly attenuated the magnitude of LTP. These data are consistent with the major contribution of vesicular mechanisms into release of glutamate (Fig.2) and ATP (Lalo et al. 2014). (C)Genetic deletion of Best1 CACC channels significantly reduced the LTP which could be rescued by application of exogenous D-Serine. These data supports the importance of the release of D-Serine from CACCs. (D)The genetic deletion of TREK-1 channels had moderate but statistically significant effect on LTP which was not rescued by the application of exogenous D-Serine. (E)The widespread (compared to (A)) inhibition of glial exocytosis in the dnSNARE mice abolished the LTP which was not rescued by exogenous D-Serine but was rescued by overcoming the deficit in the release of purinergic gliotransmitter ATP by application of its non-hydrolysable analog (ATP-γS). Note that the largest LTP out of the three transgenic lines was observed in the dn-SNARE mice.

The magnitude of LTP induced in the neocortex of Best1-KO mice under control conditions was significantly lower than in the wild-type mice but could be rescued by the application of exogenous D-Serine (Fig.5C). We also observed a deficit in the neocortical LTP induced in the TREK-1 KO mice which could not be rescued by exogenous D-Serine (Fig.5D). These data strongly support the involvement of non-vesicular glutamatergic gliotransmission, especially the release of D-Serine via Best1 channels, for the induction of long-term synaptic plasticity.

The extent of long-term potentiation of cortical fEPSPs was dramatically reduced in the dnSNARE mice (Fig.5E). Among the three transgenic lines with impaired gliotransmission, the largest deficit in the LTP was observed in the dn-SNARE mice, this could be explained by the deficit in release of both ATP and glutamate. Importantly, application of external D-serine did not rescue the LTP phenotype in the dn-SNARE mice (Fig 5E). The LTP phenotype in the dnSNARE mice was, however, rescued by application of non-hydrolysable selective agonist of P2X receptors ATPγS (Fig. 5E). These data suggest that vesicular release of ATP from astrocytes acts downstream of glutamatergic gliotransmitters in the modulation of synaptic plasticity. One of the putative mechanisms of ATP action could be lowering the threshold of LTP induction via P2X receptor-mediated down-regulation of neuronal GABAA receptors (Lalo et al., 2014a). The stronger GABA-mediated inhibition in the dnSNARE mice and, consequently, insufficient depolarization of postsynaptic neurons for activation of NMDA receptors may explain the lack of positive action of D-Serine during the LTP induction in the neocortex. The feasibility of this mechanism was corroborated by our finding that attenuation of GABA-mediated inhibition by gabazine (150 nM) facilitated the induction of LTP in the dn-SNARE mice (Fig. S4A). Another piece of evidence to support this mechanism was obtained by assessment of the contribution of NMDA receptors to synaptic potentials. The NMDAR-mediated component of fEPSPs, evaluated via the facilitatory effect of D-Serine and the inhibitory effect of D-AP5 (Fig. S4B,C) was very small under control conditions (which included physiological level of extracellular Mg^2+^). However, the contribution of NMDA receptors increased significantly when depolarization of cortical neurons was facilitated by attenuation of GABAergic inhibition by gabazine; a similar effect to the activation of ATP receptors with ATP-γS (Fig. S4C,D). In the wild-type mice, the NMDAR-mediated component was also enhanced by activation of astrocytes with TFLLR, whereas TFLLR did not have a marked effect in dnSNARE mice (Fig. S4D). These data strongly suggest that action of astrocyte-driven ATP on the postsynaptic GABA receptors (Lalo et al., 2014a) works downstream of the positive modulation of NMDA receptors by D-Serine.

### Gliotransmission plays a key role in spatial working memory and environment-induced plasticity

Combined, the results of our LTP experiments demonstrate the importance of synergism between astroglial exocytosis of ATP and glutamate and the release of D-Serine via CACCs for the regulation of synaptic plasticity. In addition to the acute synaptic plasticity directly related to learning and memory (Keck et al., 2017), astrocytes have also been implicated into slower cognitive mechanisms such as homeostatic synaptic plasticity and experience-related brain metaplasticity (Hulme et al., 2014). So, to explore a physiological role for gliotransmission in brain plasticity *in vivo*, we performed the behavioural working memory tests with the mice kept in standard housing (SH) and mice exposed to the environmental enrichment (EE). Such an approach allowed to explore the involvement of gliotransmission in neural plasticity at different time scales. We used dnSNARE mice in these *in vivo* experiments, as they are the mouse strain exhibiting the largest deficit in cortical synaptic plasticity (Fig.5). In order to evaluate the working memory we performed the spontaneous alternation test ((More et al., 2008); see also Methods). The dn-SNARE mice kept in the standard housing showed a deficit in spatial working memory in the spontaneous alternation task (Fig.6A). The percentage of correct alternations (Fig.6A) in the dn-SNARE mice was much lower as compared to their wild-type littermates; the difference was statistically significant with P < 0.005 (unpaired un-equal variance two-tailed T-test). Rather surprisingly, the exposure of mice to the environmental enrichment strongly enhanced the working memory in the dn-SNARE mice but had just a modest effect in their wild-type littermates (Fig.6A). Although some differences in the spontaneous alternation task performance between EE-exposed WT and dn-SNARE mice could be seen, it was not statistically significant.

Application of multivariate ANOVA on the number of entries, percentage of correct alternations, and percentage of correct alternations above the chance, between the factors of genotype (wild-type vs. dn-SNARE) and the treatment (SH vs. EE) revealed that the only significant difference between genotypes was within the SH condition: P = 0.002 for both correct alternations and correct alternations above chance. There was not statistically significant effect of genotype on the number of entries (P=0.91) indicating the lack of effect of dn-SNARE expression on locomotion.

The other interesting result was that only dn-SNARE mice performed significantly better upon exposure to EE (P = 0.001). The weak responsiveness of wild-type littermates to the EE might be explained by their very good performance (about 70% of correct alterations) even in standard housing conditions.

The results of our behavioral tests have two important implications. First, very good performance of the EE dn-SNARE mice in working memory test (and lack of neurological signs) strongly argue against the non-selective

“leaky” expression of dn-SNARE transgene in neurons which would have certainly led to the major deficit in synaptic signaling and memory. The lack of direct effects of dn-SNARE expression on neurons is also supported by the lack of changes in the mEPSCs (Figs.2,4). Second, the working memory deficit in the dn-SNARE mice kept in standard housing clearly demonstrated the physiological relevance of exocytotic release of gliotransmitters from astrocytes. The marked responsiveness of dn-SNARE mice to EE suggested the existence of some compensatory mechanisms which could overcome the deficit in vesicular gliotransmission. The most parsimonious explanation might be a mosaic expression of dn-SNARE across the astrocyte population and EE-induced increase in the release of gliotransmitters from non-dnSNARE astrocytes.

This hypothesis was corroborated by results of the *ex-vivo* experiments assessing the impact of EE on synaptic plasticity (Fig.6B,C) and gliotransmission (Fig.7). In line with changes in the working memory, we observed a moderate increase in the magnitude of LTP in the wild-type mice upon exposure to EE contrasting with a dramatic EE-induced increase in the dn-SNARE mice (Fig.6B). One might say that exposure to EE “restored” the LTP phenotype in the dn-SNARE mice; we observed a similar behaviour of both hippocampal and neocortical LTP (Fig. S5). An apparent weak sensitivity of wild-type mice to EE may be explained by threshold-like behaviour of LTP magnitude which is close to saturation even in the SH mice. On the other hand, very low magnitude of LTP in the dn-SNARE mice makes them very sensitive to any treatment capable of facilitating synaptic plasticity. Importantly, the EE-induced increase in the LTP in the dn-SNARE mice was significantly inhibited by the blockers of TREK-1 channels (spadin) and Best1 channels (CaCInh-A01) which had only marginal effects in the SH wild-type mice (Fig.6D, Fig.S5B). This result suggests that EE could enhance the glial signaling and increase the release of gliotransmitters, in particular via non-vesicular pathways. This notion was supported by the observation of increase in the spontaneous and NA-evoked Ca^2+^-signaling in astrocytes of EE mice (Fig.7A,B). Importantly, an increase in the astroglial Ca^2+^-signaling was un-related to the dn-SNARE expression (Fig.7A,B). In parallel with enhanced astroglial signaling, we observed the significant EE-induced increase in the release of ATP, glutamate and D-Serine from astrocytes of wild-type mice (Fig. 7C,D). The EE-induced enhancement of release of ATP and glutamate was reduced in the dn-SNARE mice as compared to wild-type littermates (Fig. 7C,D). In contrast, the EE-induced enhancement of D-Serine release was not affected.

**Figure 6.**
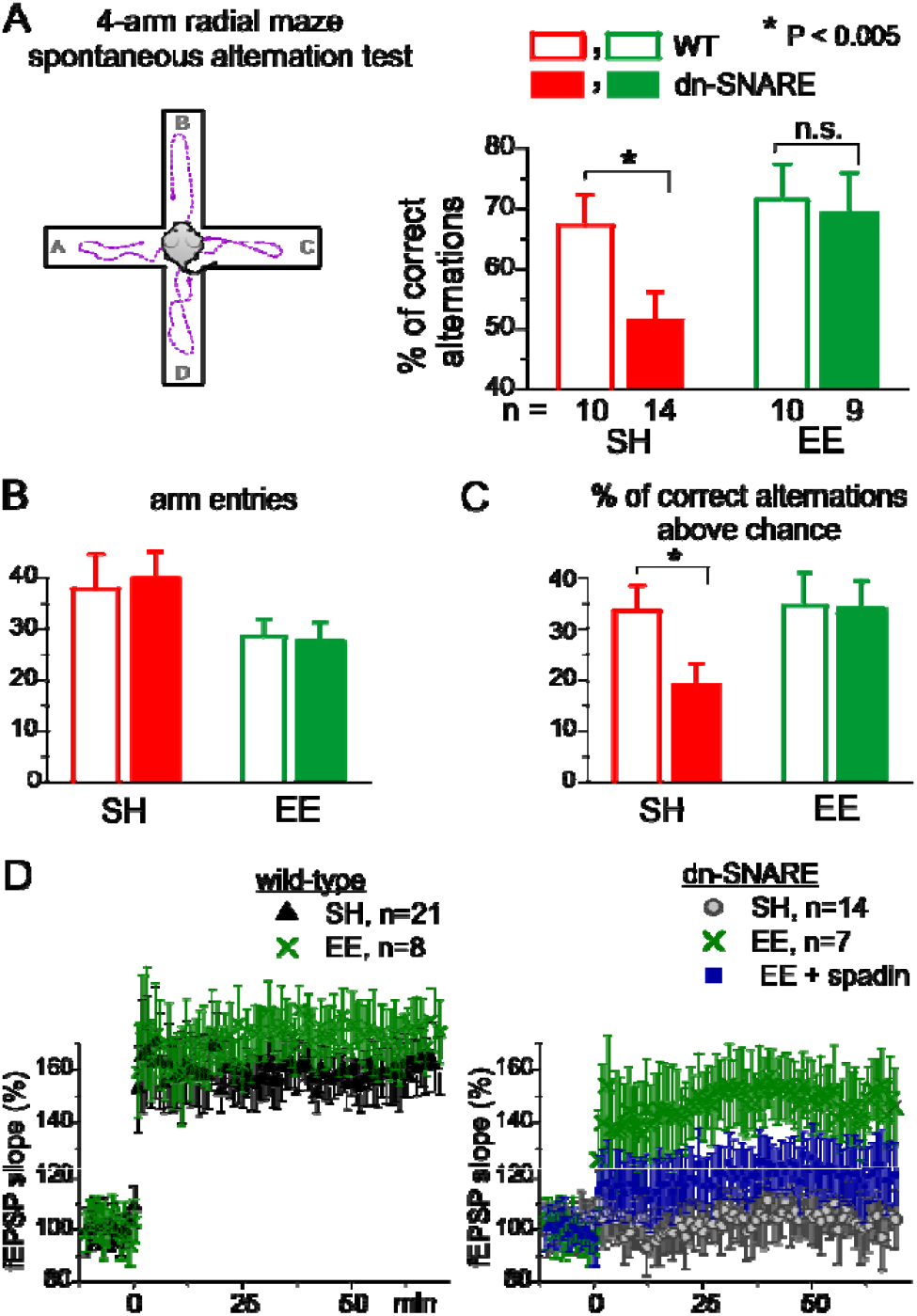
Loss of gliotransmission impairs working memory when mice are kept in standard housing. (A-C) Working memory was tested in the dn-SNARE and WT mice using Spontaneous Alternation Test: percentage of correct alternations (A), total number of entries (B), and percentage of correct alternations above the chance (C) for number of mice as indicated in (A). The dn-SNARE mice kept in standard housing (SH) show a deficit in spatial working memory whereas dn-SNARE mice exposed to the enriched environment (EE) did not show a memory deficit. (D) LTP was impaired only in the dnSNARE mice kept in standard housing (SH) but not in the enriched environment. The relative effect of inhibition of TREK-1 channels by the specific antagonist spadin was much larger in the EE mice (both WT and dnSNARE) as compared to SH-mice (compare to Fig.5D). No deficit in baseline fEPSPs was found in the dn-SNARE mice kept in standard housing or in the enriched environment.

**Figure 7.**
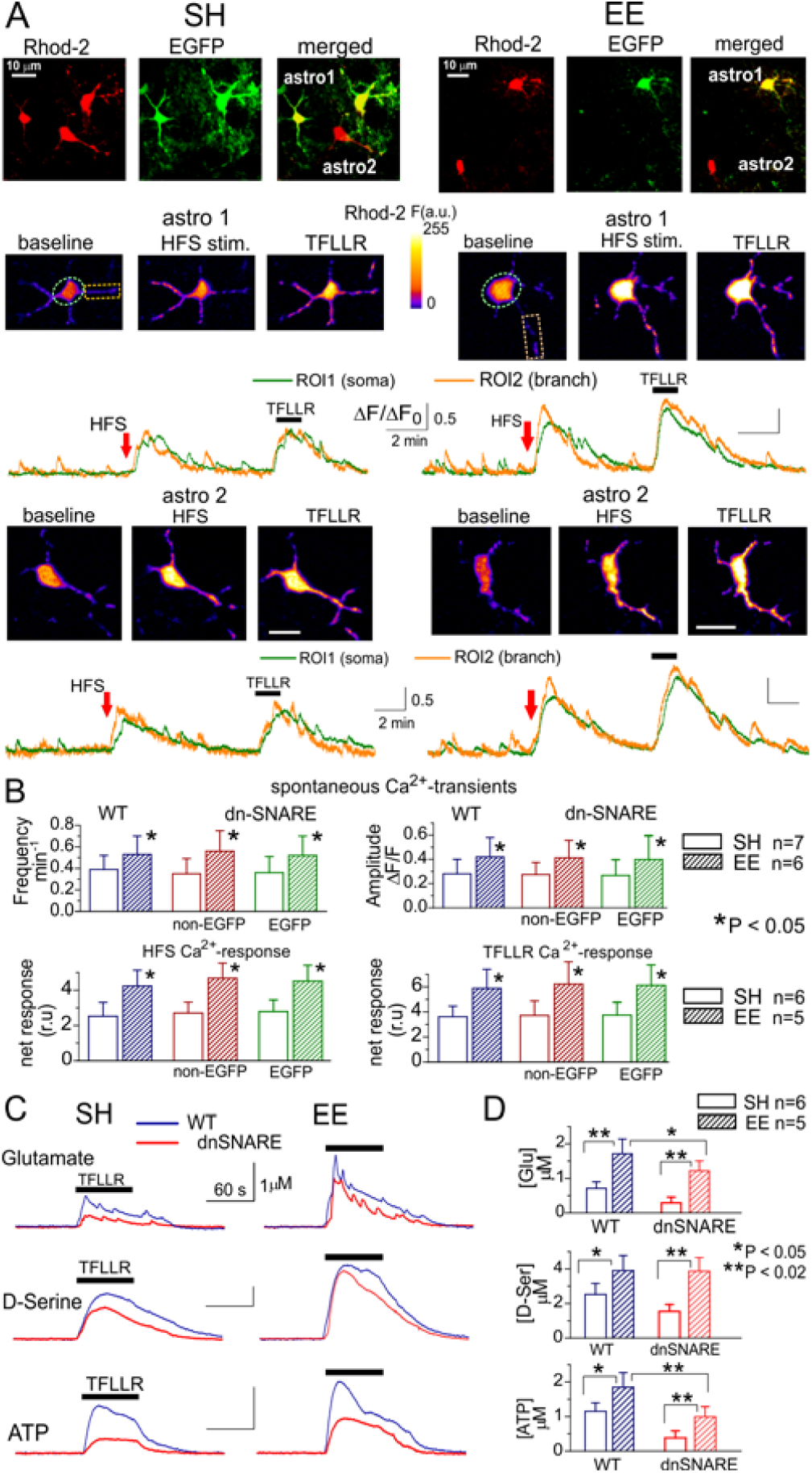
Impact of environmental enrichment on astroglial Ca^2+^-signalling and release of gliotransmitters. (A-B) Spontaneous and evoked Ca^2+^-transients were recorded in L3 neocortical astrocytes of the dnSNARE mice and their wild-type littermates kept in the standard housing (SH) and in an enriched environment (EE) using multi-photon fluorescent microscopy. (A) The representative 2-photon images of astrocytes loaded with Ca^2+^-indicator Rhod-2 and the time course of Rhod-2 fluorescence in the two highlighted ROIs before and after short burst of high frequency stimulation (HFS) of cortical afferents or direct stimulation of astrocytes with PAR-1 receptor agonist. *Left* and *Right* columns correspondingly show astrocytes from SH and EE mice. Recordings were made in the two types of astrocytes: (1) cells expressing dnSNARE-EGFP constructs and (2) unaffected “non-EGFP” astrocytes. (B) the pooled data (mean ±SD for the number of cells indicated) on the frequency and amplitude of the spontaneous Ca^2+^-transients and the net response (averaged over the whole cell image as described in Methods) to HFS and PAR-1 agonist. Note that keeping mice in an enriched environment caused a significant increase in the astrocytic Ca^2+^-signalling irrespective of dnSNARE expression. (C,D) The elevation in the extracellular concentrations of glutamate, D-Serine and ATP was triggered by activation of astrocytes with PAP-1 receptor agonist (TFLLR) and measured with microelectrode biosensors (see Methods). Release of all transmitters was significantly increased by EE both in the wild-type and dnSNARE mice. However, the release of glutamate and ATP was significantly lower in the dnSNARE mice. The EE-induced increase in the release of D-Serine was not different between genotypes. In all panels, asterisks indicate statistical significance accordingly to un-paired t-tests.

Taken together, the above results demonstrate that physiologically-attainable astroglial signaling can trigger release of glutamate via exocytosis and TREK-1 channels. Combined with Best1 channel-mediated release of D-Serine, glia-derived glutamate can activate extrasynaptic NMDARs in cortical neurons and thereby affect the induction of LTP. Our data show that synergistic action of glutamatergic gliotransmitters and astrocyte-derived ATP in regulation of synaptic plasticity can be important for learning and memory.

## Discussion

Our study provides unequivocal evidence of the physiological importance of both vesicular and channel-mediated release of glutamate and thereby can provide answers to the ongoing controversies concerning the fundamental mechanisms of gliotransmission.

Firstly, we show that neocortical L2/3 astrocytes can release glutamate by both SNARE-dependent exocytosis and non-vesicular mechanisms dependent on Best1 and TREK-1 channels (Figs 1-3). Our data suggest that exocytosis and non-vesicular release via TREK-1 channels provide major, and approximately equal, contributions into the glial release of glutamate (estimated as a charge transferred by astrocyte-derived currents in the “sniffer” cells). This estimation differs from previous “sniffer”-cell data which showed a predominance of channel-mediated release of glutamate from hippocampal astrocytes (Woo et al., 2012). This difference may, most likely, be related to the regional difference in the expression of SNARE, TREK-1 and Best1 proteins. Also, it is conceivable that the relative contributions of different pathways of gliotransmitter release can strongly depend on the metabolic state of astrocytes, e.g. its cytosolic concentration of transmitters, activity of H^+^-ATPases and vesicular transporters. One should also note the stark differences in the kinetics and Ca^2+^-sensitivity: while exocytosis is a very rapid process with relatively high (> 190 -200 nM) Ca^2+^-threshold, CaCh-mediated release is slow and has a sigmoidal dependence on cytosolic Ca^2+^ concentration with EC_50_ about 150 – 200 nM (Hartzell et al., 2008), i.e. can be partially activated even at resting Ca^2+^-level; the TREK-1 mediated release relies on modulation of K^+^ channel by G-proteins rather than Ca^2+^ (Woo et al., 2012). Hence, co-existence of exocytotic and channel-mediated release of glutamate from astrocytes can convey a great diversity and plasticity to the astrocyte-to-neuron communications, with prevalence of phasic (as in the present paper) or tonic (as in the accompanying paper by Koh et. al 2021-BIORXIV/2021/436945) activation of neuronal glutamate receptors in different brain regions or physiological context. Such variations in the spatial-temporal pattern of astroglia-derived glutamate could explain, at least partially, discrepancies in the data on the physiological effects of gliotransmission.

Secondly, we show that glia-derived glutamate can activate three types of currents in the neighbouring neurons which can be distinguished from each other and from the synaptic currents (mEPSCs) by their biophysical and pharmacological properties (Fig.2). The glia-derived currents: faster gFICs, slower gSICs and slow tonic currents (Fig. 2,4) were inhibited in different ways by perfusion of astrocytes with TeNTx and VGLUT inhibitors; they also changed differently in the dnSNARE and TREK-1 and Best1 KO mice (Fig.3). This allowed us to attribute these currents to the astroglial release of glutamate by exocytosis (gFICs) and diffusion through the TREK-1 potassium channels (gSICs) and Ca^2+^-dependent chloride channels (tonic currents). In contrast to the synaptic currents (mEPSCs), glutamatergic currents of astroglial origin are mediated predominantly by GluN2B subunit-containing NMDA receptors and exhibit slower kinetics (Fig2). Thus, astroglial-derived currents occur in the cortical neurons most likely at extra-synaptic and/or peri-synaptic *loci*, where GluN2B-containing NMDARs are predominately expressed (Paoletti et al., 2013; Papouin and Oliet, 2014). Importantly, our results revealed a dependence of all types of GluN2B-mediated glia-driven neuronal currents on glial release of D-Serine which agrees with the data of (Ferreira et al., 2017) on the preferential action of D-Serine on GluN2B-rather than GluN2A-containing NMDARs. Combined with the results of accompanying paper (Koh et al. 2021-BIORXIV/2021/436945), our data on key role of Best1 in generation of glia-driven tonic NMDAR currents suggest that this mode of astrocyte-to neuron communication can be common for many brain areas.

Finally, our data demonstrate that cortical astrocytes *in situ* can release glutamate following physiologically-relevant stimulation causing a large enhancement of glutamatergic gFICs and gSICs (Fig.2,4), which make approximately equal contributions to glia-derived activity of GluN2B-containing extrasynaptic NMDARs. Manipulations of the activity of GluN2B receptors have been reported to affect the magnitude, threshold and even the sign of subsequent plasticity (Paoletti et al., 2013). Thus, our data can link the gliotransmission to the physiological role for extrasynaptic NMDA receptors, which have been implicated into regulation of synaptic development, synaptic plasticity and metaplasticity (Paoletti et al., 2013; Papouin and Oliet, 2014).

Indeed, we have found that both vesicular and non-vesicular release of gliotransmitters is essential for the induction of NMDAR-dependent long-term potentiation. This was demonstrated by the significant reduction of the LTP magnitude upon perfusion of astrocytes with blockers of exocytosis and vesicular glutamate transporters (Fig.5A, Fig S5A) and the LTP deficit in transgenic mice with impaired release of gliotransmitters (Fig.5E, Fig.6C). Our data also indicated that glial release of both glutamate, activating extrasynaptic NMDARs (Fig.2-4, 5B), and ATP, activating extrasynaptic P2X receptors (Fig.5B, E see also (Lalo et al., 2014a)), is important for the induction of LTP. The synergism between glia-derived ATP and glutamate is produced, most likely, by the P2X receptor-mediated down-regulation of GABAergic inhibition (Lalo et al., 2014a) leading to the facilitation of NMDA receptors (Fig.S4) and up-regulation of AMPA receptors trafficking (Boue-Grabot and Pankratov, 2017). Importantly, our present (Fig.1-4) and previous (Lalo et al., 2016) observations of co-existence and co-operation between glutamatergic and purinergic gliotransmission in the neocortex go well in line with recent data obtained in hippocampus by Covelo and Araque (Covelo and Araque, 2018). Combined, these findings unveil a new level of complexity in bi-directional astrocyte-neuron communications.

Our data have provided long-awaited evidence of dnSNARE behavioural phenotype – we observed a deficit in the working memory *in vivo* that agrees with the impairment of LTP shown in the previous (Pascual et al., 2005) and current (Figs. 5,S5,6) papers. Since the working memory deficit was observed in the dnSNARE mice kept in the standard housing but not in the enriched environment, it could hardly be attributed to the allegedly “leaky” expression (Fujita et al., 2014) of dnSNARE transgene in neurons. Also, recent data support the specificity of astroglial expression of dnSNARE (Papouin et al., 2017a; Sultan et al., 2015) and verify the lack of deficit in the synaptic exocytosis of neurotransmitters (Lalo et al., 2016). Our data also argue against impact of astroglial dnSNARE expression on the channel-mediated gliotransmission (Fig.1, S1).Thus, the deficit in the LTP and working memory in dnSNARE mice can be solely attributed to the impairment of astroglial exocytosis. Yet, we would like to emphasize that we do not consider the dnSNARE mice as an ideal model to elucidate the importance of vesicular gliotransmission. Thus, in our current and previous works, the main evidences of exocytosis of glutamate and ATP have been obtained using several alternative methods (Figs 1-4; (Lalo et al., 2014a)).

Interestingly, it was one of shortcomings of dnSNARE model, namely the mosaic expression of dnSNARE in cortical areas (only in ∼50% of astrocytes), which allowed us to discover the positive effect of environmental enrichment on the release of gliotransmitters and glia-derived regulation of synaptic plasticity and memory (Figs.6,7). Furthermore, these results highlight the importance of synergism between the different pathways of gliotransmission. This synergism exists on several levels: synergism between vesicular and TREK-1 mediated release of glutamate, synergism between glial release of glutamate and D-Serine and synergism between glutamatergic and purinergic gliotransmission. These pathways can converge into threshold-like augmentation of synaptic plasticity. Inhibition of just one of the components of gliotransmission either by genetic or pharmacological manipulations can lead to impairment of synaptic plasticity and memory. On the other hand, experience and environment-related plasticity of astroglial signalling can compensate the deficit of one component (e.g. glial exocytosis) by enhance others (TREK-1 and Best1-mediated release). Our data suggest the important contribution of enhancement in astroglial adrenergic signalling in the EE-induced upregulation of release of D-serine (Fig.7A,C) which closely agrees with the results of accompanying paper (Koh et. al 2021-BIORXIV/2021/436945) on key role of astrocytic α1-adrenoreceptors in regulating of NMDAR-tone via Best1 and implications of this cascade into synaptic metaplasticity. The synergism between different pathways of gliotransmission, which can undergo experience-dependent changes, could explain the ongoing controversies about the impact of manipulations of astrocytic signalling on synaptic plasticity (Bazargani and Attwell, 2016; Papouin et al., 2017b; Savtchouk and Volterra, 2018).

Our data on the importance of Ca^2+^-dependent channel-mediated astrocytic release of D-Serine for the generation of glia-driven extrasynaptic gFICs and gSICs and modulation of synaptic plasticity (Fig.3,5,7,S5) and the data on role for astroglia-derived D-Serine in cognitive flexibility (accompanying paper Koh et. al 2021-BIORXIV/2021/436945) could resolve the emerging controversy about physiological relevance of glial as source of D-Serine. Despite the large body of evidence supporting the importance of glial D-Serine (Henneberger et al., 2010; Panatier et al., 2006; Sultan et al., 2015), the ability of astrocytes to release D-Serine has been recently questioned (Wolosker et al., 2016). One should note that latter work relied mainly on the immunostaining of neurons and astrocytes to with antibodies to serine racemase and D-Serine whose specificity remains questionable (Papouin et al., 2017b). In contrast, the present and accompanying papers (Koh et. al 2021-BIORXIV/2021/436945)

One should emphasize that even the studies questioning astroglia-specific expression of serine racemase, support the notion of significant (if not predominant) accumulation of D-Serine in the astrocytes. Hence, the relative contribution of glial and neuronal sources for the release of D-Serine will depend mainly on the efficiency of the particular mechanisms of release. So far, there is no direct evidence that neurons possess an efficient mechanism for the activity-dependent release of D-serine. Wolosker and Coyle (Wolosker et al., 2016) suggested the Asc-1 or other transporters act as the main pathway. In comparison to Best1 channel, allowing the movement of multiple molecules, transporters can only release a single molecule per single conformational change and are therefore intrinsically slow. Even Na^+^-dependent transporter-mediated release cannot be activated as fast as Ca^2+^-dependent Best1 channels. So, one could hardly expect a predominant role of neurons in the activity-dependent release of D-Serine. Indeed, our experiments in isolated neurons and neurons in slices, which allowed us to untangle glial and neuronal sources of D-Serine (Fig.S3), showed that neuronal release on its own could not provide enough D-Serine to maintain the activity of synaptic and extra-synaptic NMDARs. Furthermore, the whole complex of our data presented here (Fig.2,3,5,7,S3) is in a good agreement with data on the important physiological role of astroglial release of D-Serine (Panatier et al., 2006; Papouin et al., 2017b; Sultan et al., 2015).

**To conclude**, our results demonstrate that release of gliotransmitters from astrocytes under physiological conditions involves both vesicular and non-vesicular mechanisms and synergism between these mechanisms is essential for glia-driven regulation of synaptic transmission and plasticity

## ACKNOWLEDGEMENTS

This work was supported by grant from BBSRC UK BB/K009192/1 to Y.P. and BB/N012941/1 to Y.P. and C.J.L.. We thank Prof. A. Volterra and Prof. F. Kirchoff for early discussion and Prof. P.G. Haydon for early discussion and providing of dnSNARE mice.

## AUTHOR CONTRIBUTIONS

Conceptualization, U.L., C.J.L., and Y.P.; Methodology, L.M., J.M, M.W., C.J.L., and Y.P.; Investigation, U.L., S.R-N., A.B., W.K., L.M., J.M., M.W., and Y.P.; Writing – Original Draft, U.L., J.M., and Y.P.; Writing – Review & Editing, M.W., C.J.L., and Y.P.; Formal Analysis, U.L., L.M., K.W., and Y.P.; Resources, J.M., M.W., and C.J.L.; Funding Acquisition, C.J.L., and Y.P; Supervision, J.M, M.W., C.J.L., and Y.P.

## STAR METHODS

## CONTACT FOR REAGENTS AND RESOURCES SHARING

Further information and requests for resources and reagents should be directed to and will be fulfilled by the Lead Contact, Yuriy Pankratov

## EXPERIMENTAL MODEL AND SUBJECT DETAILS

### Animals, genotyping and housing

All animal work has been carried out in accordance with UK legislation (ASPA) and “3R” strategy; all experimental protocols were approved by University of Warwick Ethical Review Committee and Animal Welfare Committee. Experiments were performed in 2-5 months old C57BL/6 mice and transgenic mice with knockout of TREK-1 potassium channels (Namiranian et al., 2011) (TREK-1 KO), Best1 channels (Marmorstein et al., 2006) (Best1 KO) and dnSNARE transgenic mice with impairment of astroglial exocytosis (Lalo et al., 2014a; Papouin et al., 2017a; Pascual et al., 2005). Genotypes of all animals were verified by PCR from the ear samples.

The dnSNARE mice originated from the Prof. P.G. Haydon’s lab (Department of Neuroscience, Tufts University School of Medicine, Boston, MA 02111) and were bred and genotyped as previously described (Pascual et al., 2005; Sultan et al., 2015). We would like to emphasize that absence of neuronal expression of dnSNARE transgene in this particular line has been verified in several recent studies (Papouin et al., 2017a; Sultan et al., 2015). Administration of Dox to dnSNARE mice (Lalo et al., 2014a; Pascual et al., 2005) has been removed 4 weeks prior to recordings.

Data obtained in the C57BL/6 mice did not differ significantly from data obtained in the littermate non-carriers of the TREK-1 KO, Best1 KO and dn-SNARE transgenes of the same age (n=5-8 for each type of experiments). For clarity, all data referred here as wild-type were reported for C57BL/6 mice.

All animals were kept on a 12-12 light-dark cycle with water and food *ad libitum*. Standard housing conditions (SH) included a bedding and a cardboard tube; mice were housed in groups of 2-3 siblings. Enriched environment housing (EE) was provided via a large cage (Ferplast Duna Hamster Cage: 39 cm w x 55 cm l x 27 cm h) containing bedding, running wheel and toys (ladders, rope, shelters, tunnels) arranged on two levels. The cage top was covered with high-ceiling lid so that mice could also explore their environment vertically. To provide novelty, toys were moved around and new toys introduced every few days. Mice were housed in larger groups (6 – 10 mice) from birth to facilitate social interactions. At weaning (3 weeks) females were removed and after then males and females were housed separately in the groups of 3-8 mice.

### HEK293 cell cultures and transfections

To generate glutamate-sensing HEK293 cells, the pCI_neo_-GluRI-Y497L plasmid (Woo et al., 2012) was used. The full-length GluR1-Y497L sequence was amplified with primers providing XhoI and EcoRI restriction sites by standard PCR techniques. The resulting fragment was then sub-cloned into the corresponding sites of the pIRES-EGFP vector to enable the isocistronic expression of GluR1-Y497L with EGFP to aid the unambiguous identification of transfected cells. HEK293 cells were cultured in DMEM supplemented with 10% FBS (Invitrogen, Carlsbad, CA, USA). Cells were transfected by the calcium phosphate method as described (Canal et al., 2011) and analysed 48 hours after transfection.

## METHOD DETAILS

### Slice and cell preparation

Mice were anaesthetized by halothane and then decapitated, in accordance with UK legislation. Brains were removed rapidly after decapitation and placed into ice-cold physiological saline containing (mM): NaCl 130, KCl 3, CaCl_2_ 0.5, MgCl_2_ 2.5, NaH_2_PO_4_ 1, NaHCO_3_ 25, glucose 15, pH of 7.4 gassed with 95% O_2_ - 5% CO_2_. Transverse slices (280 μm) were cut at 4° C and then placed in physiological saline containing (mM): NaCl 130, KCl 3, CaCl_2_ 2.5, MgCl_2_ 1, NaH_2_PO_4_ 1, NaHCO_3_ 22, glucose 15, pH of 7.4 and kept for 1 - 4 h prior the cell isolation and recording.

Neocortical and hippocampal astrocytes were acutely isolated using the modified “vibrating ball” technique (Lalo et al., 2014a; Lalo and Pankratov, 2017; Lalo et al., 2006). The glass ball (200 µm diameter) was moved slowly some 10 - 50 µm above the slice surface, while vibrating at 100 Hz (lateral displacements 20 - 30 µm). The composition of external solution for all isolated cell experiments was (mM): 135 NaCl; 2.7 KCl; 2.5 CaCl_2_; 1 MgCl_2_; 10 HEPES, 1 NaH_2_PO_4,_ 15 glucose, pH adjusted with NaOH to 7.3. This technique preserves the function of membrane proteins and therefore is devoid of many artefacts of enzymatic cell isolation and culturing procedures. In particular, vibro-dissociated astrocytes retain many morphological features (e.g. GFAP-EGFP fluorescence, size, proximal processes) and functional properties (e.g. high potassium conductance, glutamate transporters, Ca^2+^-signalling) whilst being completely isolated from neuronal somata and nerve terminals.

### Fluorescent microscopy and Ca^2+^-imaging

Two-photon imaging of astrocytes and neurones was performed using Zeiss LSM-7MP multi-photon microscope coupled to the SpectraPhysics MaiTai pulsing laser; experiments were controlled by ZEN LSM software (Carl Zeiss, Germany). Images were further analyzed off-line using ZEN LSM (Carl Zeiss) and ImageJ (NIH) software. To monitor the cytoplasmic free Ca^2+^concentraton ([Ca^2+^]_i_) in situ, astrocytes of neocortical slices were loaded via 30 min incubation with 1 µM of Rhod-2AM (dn-SNARE mice) or Calcium Green-2AM and sulphorhodamine 101 (other mice) at 33°C. Alternatively, individual astrocytes were perfused with intracellular saline containing 50 µM of Calcium Green-2 potassium salt; all dyes were from *MolecularProbes/Invitrogen*. Fluorescence was excited at 810 nm; Calcium Green-2 signal was observed at 520 ± 10 nm, Rhod-2 signal was observed at 590 ± 20 nm. The [Ca^2+^]_i_ levels were expressed as ΔF/F ratio averaged over a region of interest (ROI). For analysis of spontaneous Ca^2+^–transients in astrocytes, at least 3 ROIs located over dendrites and 1 ROI located over the soma were chosen. Overall Ca^2+^ -response to agonists of astroglial receptors or synaptic stimulation was quantified using an ROI covering the whole cell image.

Prior to recordings (both in acutely isolated cells and *in situ*), astrocytes were identified by their morphology under DIC observation and fluorescence (EGFP in astrocytes from dn-SNARE mice and loading with Rhod-2 or SR- 101); in some experiments, astrocytes were immunolabeled with antibodies to astroglia-specific markers (GLT1, GFAP). Identification of astrocytes was confirmed at the end of experiments by electrophysiological characterization as described previously (Lalo et al., 2014a; Lalo et al., 2006).

### Immunostaining and co-localisation analysis

For immunolabelling of vesicular transporters, secretory organelles and astroglial markers, acutely isolated astrocytes were incubated with 0.1 µg/ml of following antibodies: mouse monoclonal anti-VGLUT1 (McKA1), mouse monoclonal anti-GLT-1 (10B7, Abcam); mouse monoclonal anti-NeuN (A60), rabbit monoclonal anti-S100b (EP1576Y, Millipore); goat monoclonal anti-SV2A, mouse monoclonal anti-NG2, rabbit polyclonal anti-GFAP (Sigma). Prior to cell loading, antibodies were conjugated to the green fluorescent dye Atto488 (GLT-1, GFAP, S100b) or red fluorescent dye Atto594 (NeuN, VGLUT-1, SV2) using Lighting-Link antibody conjugation system (Innova Bioscience, Cambridge, UK) accordingly to the manufacturer’s protocol. Antibodies to VGLUT1, were applied to living astrocytes directly; other antibodies were conjugated with BioPORTER protein delivery reagent (Genlantis, San Diego, CA) 10 min prior to incubation. Immediately after isolation from the brain slice, living astrocytes were pre-incubated with 2% of normal bovine serum (Sigma) in the extracellular saline for 20 min to block unspecific binding sites. After then, cells were gently washed two times with clean extracellular saline for 5 min and then incubated with antibodies for 40 min at room temperature. After incubation, cells were washed with laminar flow of extracellular solution in the microscope recording chamber for 15 min prior to image recording. Fluorescence was excited at 820 nm; ATTO-488 signal was observed at 520 ± 10 nm, ATTO-594 signal was observed at 590 ± 20 nm. Co-localisation analysis of images was carried out using ImageJ software as described previously (Lalo et al., 2014a; Lalo et al., 2016).

### Electrophysiological recordings

Whole-cell voltage clamp recordings from cortical neurones and astrocytes and HEK293-GluR1-L497Y cells were made with patch pipettes (4 - 5 MΩ for neurons and HEK293 cells and 6-8 MΩ for astrocytes) filled with intracellular solution (in mM): 110 CsCl (KCl for astrocytes), 10 NaCl, 10 HEPES, 5 MgATP, 0.1 EGTA, pH 7.35; Currents were monitored using an MultiClamp 700B patch-clamp amplifier (Axon Instruments, USA) filtered at 2 kHz and digitized at 4 kHz. Experiments were controlled by Digidata1440A data acquisition board (Axon Instruments, USA) and WinWCP software (Strathclyde University, UK); data were analyzed by self-designed software. Liquid junction potentials were compensated with the patch-clamp amplifier. The series and input resistances were respectively 5 -7 MΩ and 500-1100 MΩ in the HEK293 cells and neurons and 8-12 MΩ and 30 - 140 MΩ in the astrocytes; both series and input resistance varied by less than 20% in the cells accepted for analysis. For activation of synaptic inputs, axons originating from layer IV-VI neurones were stimulated with a bipolar coaxial electrode (WPI, USA) placed in the layer V close to the layer IV border, approximately opposite the site of recording; stimulus duration was 300 µs. If not stated otherwise, the stimulus magnitude was set 3 - 4 times higher than minimal stimulus (typically 0.8–1.2 μA) adjusted to activate the single-axon response in the layer II pyramidal neurones as previously described (Lalo et al., 2014a; Lalo et al., 2016). For induction of long-term plasticity in the neocortex, 5 episodes of theta-burst stimulation (TBS) were used; each TBS episode consisted of 5 pulses of 100 Hz stimulation, repeated 10 times with 200 ms interval (total 50 pulses per episode). For induction of LTP in the hippocampal CA1 area (Fig.S5) a single 1 s-long train of 100 Hz stimulation was used.

### Detection of glutamate release from astrocytes using “sniffer”-cells

After isolation from brain slice, neocortical or hippocampal astrocytes were incubated with 1μM Ca^2+^-indicator Calcium Green-2AM and astroglial marker SR101 for 30 min, re-suspended in small volume (200-300 μL) of fresh extracellular medium and placed over cultured HEK293 cells expressing mutant GluA1-L497Y receptor (Woo et al., 2012); the mutant receptor has much higher affinity to glutamate and desensitizes much slow than wild-type GluA1 receptors.

To evaluate release of glutamate, the transmembrane currents were recorded in the HEK293-GluA1-L497Y cells which had an astrocyte lying on their surface; simultaneously, astrocytes were activated by fast application of agonists of PAR-1 receptors (10 μM TFLLR), α1-adrenoreceptors receptors (1 μM noradrenaline) or endocannabinoid CB1 receptor agonist anandamide (AEA, 250 µM). These agonists have been demonstrated to induce the robust Ca^2+^-elevation and activate G-proteins in the astrocytes (Lalo et al., 2014a; Pankratov and Lalo, 2015; Rasooli-Nejad et al., 2014; Woo et al., 2012) but did not cause any response in HEK293-GluA1-L497Y or plain HEK293 cells when applied without astrocytes placed over them.

As it was previously shown (Lalo et al., 2014a; Lalo et al., 2006), the spatial density of acutely isolated cells in our experiments was rather low so we could easily select HEK cells contacting astrocytes with no other cell lying in the immediate vicinity (e.g. as shown in Figures 1 and S1). We performed recordings from the HEK293 cell-astrocyte couples, which were separated from any other cell at least by 15-20 microns of free space to ensure that HEK293-GluA1-L497Y cells were activated by glutamate released only from contacting astrocyte. Identification of isolated astrocytes was confirmed at the end of recordings by electrophysiological characterization as described previously (Lalo et al., 2014a; Lalo et al., 2006). In some experiments, morphological and electrophysiological identification of astrocytes was also verified by labeling with antibodies to astroglial (Fig.1, Fig.S1) and neuronal markers as described above; the absence of neuronal material on acutely isolated astrocytes was also shown previously in (Lalo et al., 2014a).

## QUANTIFICATION AND STATISTICAL ANALYSIS

All culture and animal experiments were quantified blinded as to the experimental manipulation. Mice were randomly allocated to experimental groups and all data collected in the course of these studies were included in the individual analyses. In the whole-cell electrophysiological recordings, data obtained in cells where series or input resistance varied more than 20% in course of experiment were excluded from further analysis.

All data are presented as mean ± SD, the statistical significance of difference between data groups was tested by two-population t-test, unless indicated otherwise. The spontaneous transmembrane currents recorded in neurones were analysed off-line using methods described previously (Canal et al., 2011; Lalo et al., 2014a; Lalo et al., 2016; Pankratov et al., 2007).

The amplitude and kinetics of synaptic currents were determined by fitting with one or two (where appropriate) theoretical waveforms with mono-exponential rise and decay phases. As a rule, mean square error of fit amounted to 5 -20% of peak amplitude depending on the background noise.

The amplitude distributions of spontaneous and evoked currents were analyzed with the aid of probability density functions and likelihood maximisation techniques; all histograms shown were calculated as probability density functions. The amplitude distributions were fitted with either multi-quantal binomial model or bi-modal function consisting of two Gaussians with variable peak location, width and amplitude. The decay time distributions were fitted with bi-modal functions. Parameters of models were fit using likelihood maximisation routine. To monitor and analyse the time course of changes in the amplitude and frequency of spontaneous currents, the amplitude and frequency were averaged over the 1 min time window.

### Spontaneous Alternation test for spatial working memory

A classical 8-radial arm maze for mice (More et al., 2008; More et al., 2007) was placed on the water tank with 4 out 8 arms kept open to form a cross. Each mouse was singly released in the centre of the maze and tracked for 10 minutes with *AnyMaze4*.*99* tracking software. The sequence and the number of arm entries were scored. A correct alternation was considered when a mouse made only one repetition in 5 entries. Prior to working memory test, mice were assessed by a battery of neurological tests (Wolf et al., 1996); all testing was run blind for genotype. All mice had neither neurological signs nor infections.

### Measurement of extracellular concentration of ATP, Glutamate and D-Serine in the brain tissue

The concentration of ATP, Glutamate and D-Serine within cortical slices was measured using microelectrode biosensors obtained from Sarissa Biomedical Ltd (Coventry, UK). These biosensors consist of platinum electrodes coated with D-serine degrading enzyme (DAAO) embedded in a perm-selective layer) and used accordingly manufacturer’s instructions (http://www.sarissa-biomedical.com/products/sarissaprobes.aspx). A detailed description of the properties of biosensors and recording procedure has been published previously in (Gourine et al., 2010; Lalo et al., 2014a; Pankratov and Lalo, 2015; Papouin et al., 2017a; Rasooli-Nejad et al., 2014). Briefly, biosensors consisted of specific transmitter-metabolizing enzymes immobilized within a matrix on thin (25-50 µM) Pt/Ir wire. This allowed insertion of the sensors into the cortical slice and minimised the influence of a layer of dead surface tissue. The concentrations of ATP, glutamate or D-Serine have been calculated from difference in the signals of two sensors: a screened ATP/Glu/D-Ser-sensor and screened null-sensor, possessing the matrix but no enzymes. This allowed to compensate for release of any non-specific electro-active interferents. Biosensors show a linear response to increasing concentration of transmitters and have a rise time less than 10 s. Biosensors were calibrated with known concentrations (10 μM) of corresponding transmitter (ATP, Glu or D-Ser) before the slice was present in the perfusion chamber and after the slice had been removed. This allowed compensation of any reduction in sensitivity during the experiment. Biosensor signals were acquired at 1 kHz with a 1400 CED interface and analyzed using Spike 6.1 software (Cambridge Electronics Design, Cambridge, UK).

