## Supplemental Figures for "Synergy between vesicular and non-vesicular gliotransmission regulates synaptic plasticity and working memory"

**Supplementary Figures**

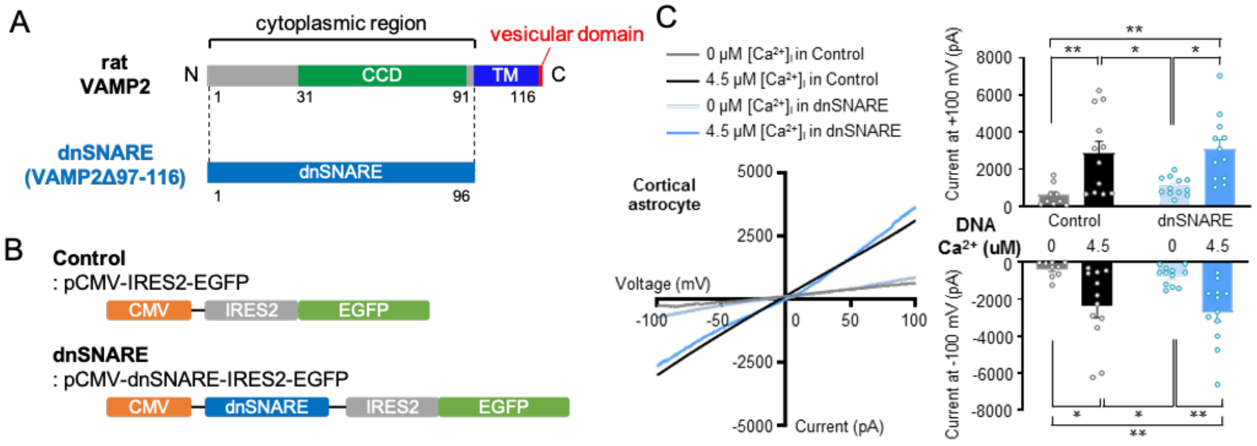


**Figure S1. Impact of dnSNARE expression on CACC current with in cortical astrocytes**
The primary cultured neocortical astrocytes were electroporetically transfected with dnSNARE and control plasmids and the calcium activated chloride currents (CACC) were evaluated as previously described (Woo et al., 2012). The CACCs are mediated predominantly by the Best1 channels, as demonstrated in Woo et al., 2012.
**(A, B)** Construction of dnSNARE expressing plasmids. (C) Left: the representative I-V relationship of CACC current in the control and dnSNARE-expressing astrocytes absence and presence of intracellular $Ca^{2+}$.
Right: the pooled data on the CACC amplitude at -100 mV and +100 mV in each condition. Data are shown as mean±SEM for 9 cells at 0 $Ca^{2+}$ and 12 cells for 4.5 mM $Ca^{2+}$, dots show individual cells. Asterisks indicate the statistical significance of difference (unpaired t-test): *p< 0.05, **p<0.01, ***p<0.001. The difference between CACC in the control and dnSNARE-expressing astrocytes was not significant both at 0 and 4.5 mM $Ca^{2+}$ (statistical power > 0.9).
This result demonstrate that constitutive surface expression of Best1 is not affected by the expression of dnSNARE in the cortical astrocytes.

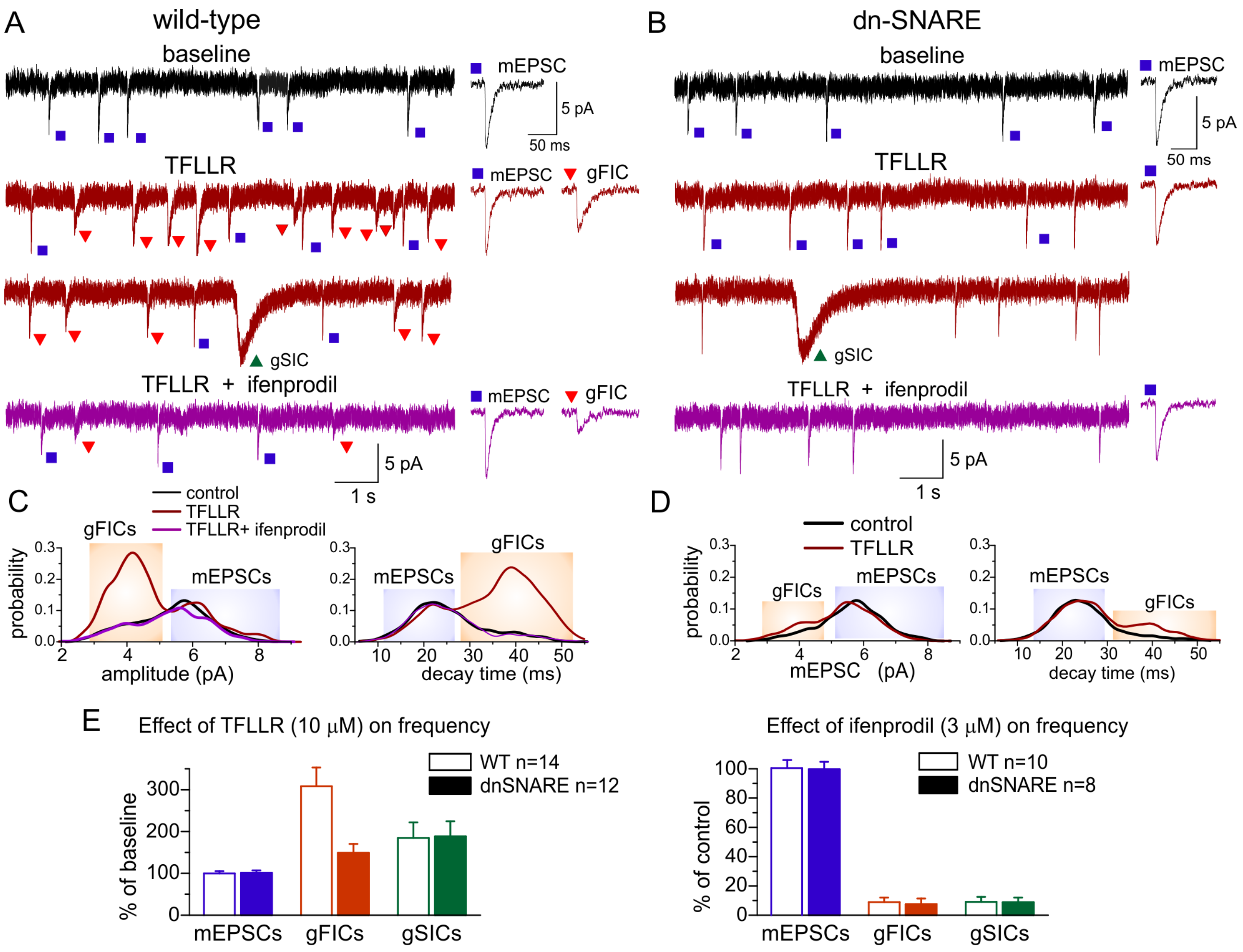


**Figure S2. Comparison of mEPSCs and glia-derived transient currents in the wild-type and dnSNARE mice.** (A-D) Whole-cell currents were recorded in pyramidal neurons of layer 2/3 of neocortical slices from wild-type (A,C) and dnSNARE (B,D) mice at -40 mV in the presence of picrotoxin, TTX, DNQX, PPADS and 5-BDBD. Activation of astrocytic $Ca^{2+}$signalling by PAR1 receptor agonist TFLLR (Lalo et al. 2014; Woo et al.2012) elicited the burst of transient NMDAR-mediated currents, similar to NEP-induced bursts shown in the Fig.2. The TFLLR-elicited burst was occulted by application of selective GluN2B-inhibitor ifenprodil.

(A,B) The representative whole-cell currents recorded in the neurons of WT and dnSNARE mice before (baseline) and during application of TFLLR alone and in the presence of ifenprodil (TFLLR was applied for 5 min with 15 min interval). Dots indicated the transient NMDAR-mediated currents separated accordingly the decay time and amplitude as shown in the corresponding distributions below (C,D).

The transient currents was subdivided into three groups: *(1) mEPSCs* – events with decay time decay time around 20 ms and larger quantal size (right peak at the amplitude distribution and left peak at the decay time distribution), the frequency of these currents was not strongly affected by stimulation of astrocytes with TFLLR;. *(2) gFICs* - currents with decay time around 40-45 ms and smaller quantal size (left peak at the amplitude and right peak at the decay time distributions), their significantly increased after stimulation of astrocytes in the wild-type but not in the dnSNARE mice; these currents were denoted as glia-induced faster currents (gFICs).

(3) gSICs – events with decay time > 100 ms (distribution shown in Fig.2C) , their frequency significantly increased after stimulation of astrocytes both in the WT and dnSNARE mice; these currents were denoted as glia-induced slower currents (gSICs). The average waveforms (20 event each) of mEPSCs and gFICs are shown in the insets in the panels (A,B).

(E) Pooled data (mean±SD) on the increase in the frequency of three types of transient currents before (baseline) and after application of TFLLR and changes in the frequency after application ifenprodil in the presence of TFLLR (compared to application of TFLLR alone). Note that only gFICs and gSICs were affected by TFLLR and ifenprodil. Effect of TFLLR on frequency of gFICs and gSICs was stastically significant with $P < 0.005$ (paired t-test) both in the wild-type and dnSNARE mice. Only gFICs were sensitive to the astrocytic dnSNARE expression, the difference in the effect of TFLLR on gFICs frequency was significant with $P < 0.005$ (unpaired t-test). These data verify the separation of transient currents into mEPSCs of neuronal origin and glia-derived gFICs and gSICs. The data strongly support the origin of gFICs from vesicular release of glutamate from astrocytes. The lack of changes in the mEPSCs in the dnSNARE mice argues against a “leaky” transgene expression in neurons.

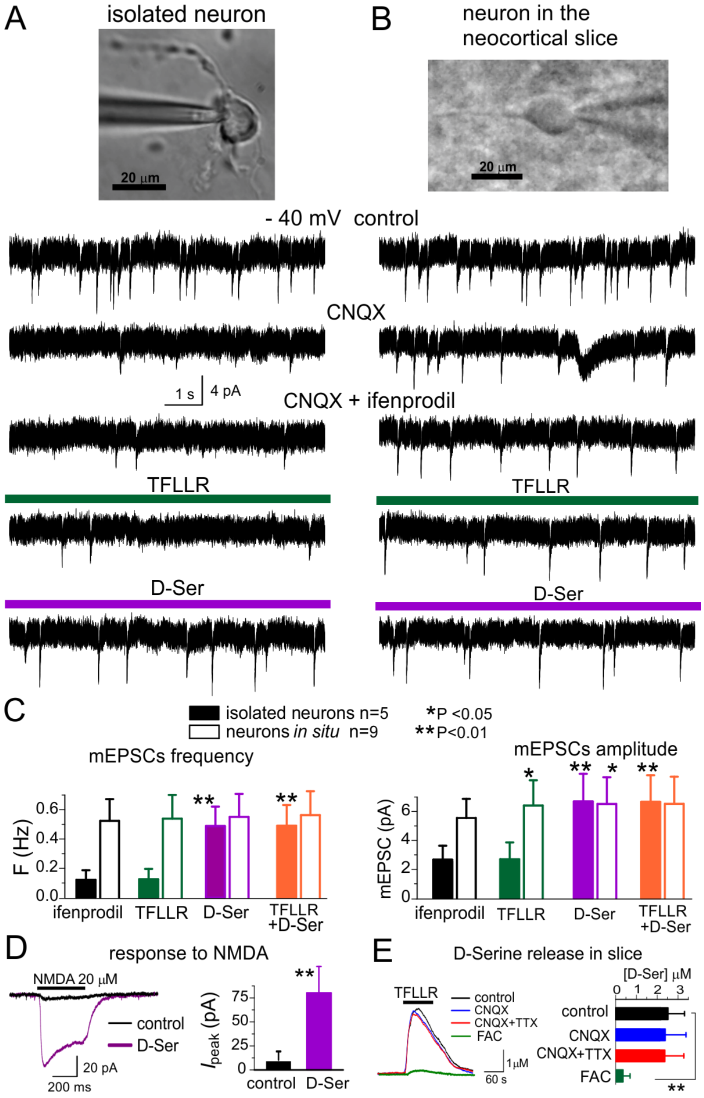


**Figure S3. The lack of efficient release of D-Serine from neurons.**

(A) Modulation of NMDA receptors by D-Serine was evaluated in acutely isolated neocortical neurons retaining the functional glutamatergic synapses (Lalo et al. 2016) and compared to the pyramidal neurons present in the neocortical slices (B).

(A,B) Representative whole-cell currents recorded at -40 mV in the presence of picrotoxin, TTX , PPADS and 5-BDBD (control) before and after consecutive application of CNQX, ifenprodil, glial activator TFLLR and exogenous D-Serine (after washout of TFLLR). The transient events which can be seen in the isolated neuron (A) in the control are miniature excitatory synaptic currents, as demonstrated in Lalo et al. 2016. Note that kinetics, amplitude and frequency of these events recorded in the control is very close to mEPSCs recorded in slice (B). Inhibition of AMPA receptors with CNQX markedly suppressed the spontaneous activity in the isolated neuron as compared to its counterpart in slice (B). The CNQX –insensitive currents were completely eliminated by application D-APV both in the isolated neuron and slice (data are not shown) confirming they were mediated by NMDARs. The spontaneous NMDAR currents recorded in presence of ifenprodil from the neuron in slice consisted predominantly of mEPSCs of neuronal origin (as shown in Fig.2S). Note the lack of the effect of ifenprodil on the isolated neuron. The amplitude and frequency of NMDAR-mediated mEPSCs in the isolated neurons were much lower in comparison to the neuron in slice. However, spontaneous currents in the isolated neurons were restored in the presence of exogenous D-Serine (10 μM). The most parsimonious explanation is that most NMDARs on the membrane of isolated neurons were not exposed to sufficient concentrations of co-agonist until exogenous D-Serine was applied.

(C) The pooled data (mean±SD) on the amplitude and frequency of the NMDAR mEPSCs recorded under different conditions. Asterisks indicate the statistical significance of the effect of application of TFLLR and/or D-Serine as compared to the mEPSCs under ifenprodil. Note the striking difference in the effect of exogenous D-Serine (10 μM): the amplitude and frequency of NMDAR-mediated currents in the isolated neurons dramatically increased whereas frequency of mEPSCs recorded in slices did not undergo significant changes.

Application of agonist of PAR-1 receptor TFLLR did not have any effect on the frequency of ifenprodil-insensitive mEPSCs both in the isolated neuron and in slice. The amplitude of mEPSCs recorded in the neuron *in situ* was moderately increased by TFLLR, the similar effect was achieved by exogenous D-Serine; combining the TFLLR and D-Serine did not cause any further increase in the amplitude. These results suggest that 1) neurons in slices are exposed to much higher concentration of NMDAR co-agonists than isolated neurons devoid of glial influence; 2) activation of astrocytes with TFLLR cause further increase in the D-Serine level which, most likely, saturates the co-agonist binding site of NMDARs. The data shown in panels (D) and (E) corroborate this conclusion.

(D) The whole-cell currents elicited in the isolated neurons by fast application of NMDA (Pankratov et al. 2002; Palygin et al. 2016) were recorded at membrane potential of -40 mV in the control and in presence of D-Serine. The significant increase in the amplitude of the response under exogenous D-Serine supports the lack of basal neuronal release of NMDAR co-agonist.

(E) The extracellular concentration of D-Serine in the neocortical slice was measured with the aid of microelectrode biosensor (Rasooli-Nejad et al. 2014). Left graph shows examples of TFLLR-induced elevation of D-Serine level; the right graph shows mean±SD for peak D-Serine concentrations (n=8 for control and n=5 for other conditions. Inhibition of neuronal activity with CNQX and/or TTX did not affect the TFLLR-induced release whereas pre-incubation of slice with glial metabolic poison fluoroacetate (20 min, 3 mM) significantly inhibited the release suggesting the astroglia as a main source of D-Serine.

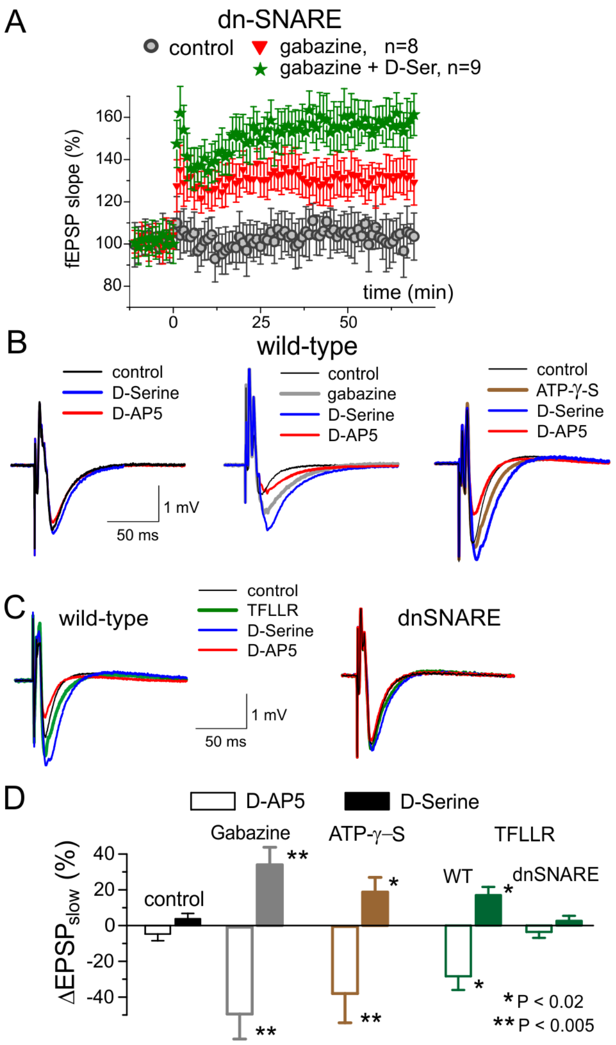


**Figure S4. The down-regulation of GABA receptorsby astrocyte-derived ATP facilitates the activation of NMDA receptors.**

(A) GABA receptor antagonist gabazine (150 nM) significantly increased the magnitude of LTP in dn-SNARE mice. In the presence of gabazine, D-Serine became capable of increasing the LTP magnitude in the neocortex of dnSNARE mice. These data suggest that down-regulation of inhibitory receptors by ATP released from astrocytes is essential for the induction of LTP in the neocortex.

(B,C) Representative field EPSPs (average of 10) evoked in the layer 2/3 of somatosensory cortex by single stimulus. Agonist (D-Serine, 10 $\mu$M) and antagonist (D-APV, 30 $\mu$M) of NMDA receptors were applied in baseline conditions (B, left) and in presence of 150 nM gabazine (B,middle), 10 $\mu$M ATPγ-S (B, right) and PAR-1 agonist (C).

(D) The pooled data (mean±SD, n=7 in each case) for the effects of D-Serine and D-APV on amplitude of slow component of fEPSP, averaged over 5 ms time period immediately after the peak.

Effects of NMDA receptors modulators were small in control but increased after suppression of GABA receptors either by gabazine or by application of ATPγ-S or TFLLR (as shown in Figure 5). Note the lack of effect of TFLLR on the NMDA component of fEPSPs in the dnSNARE mice.

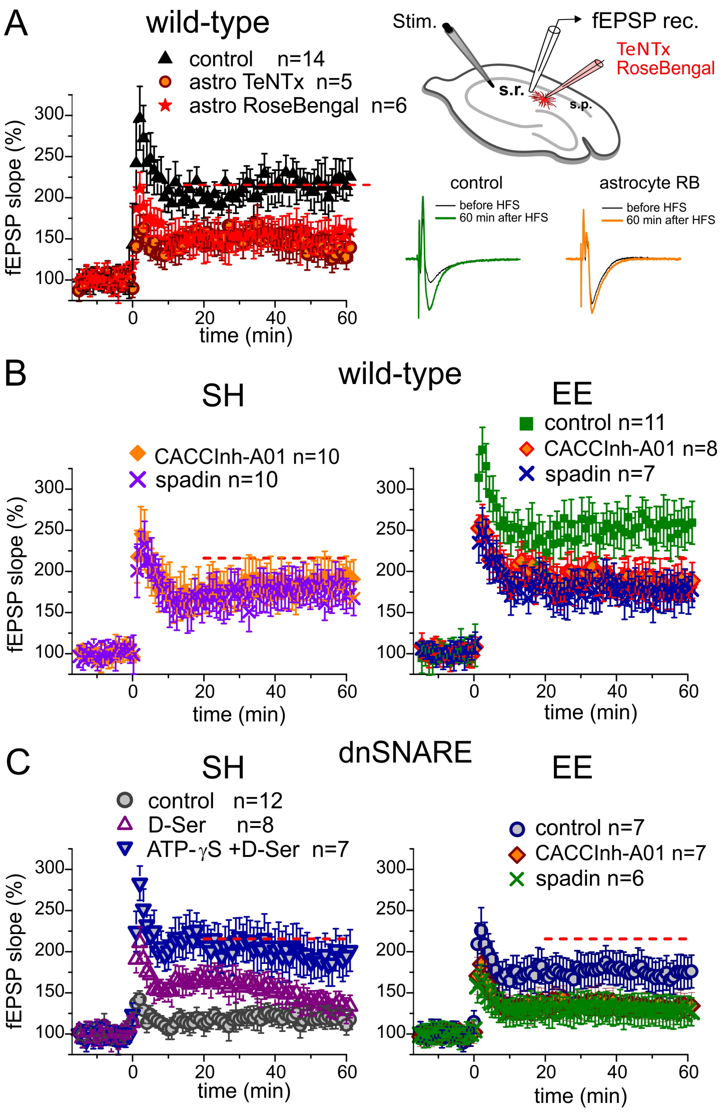


**Figure S5. Release of gliotransmitters is essential for regulation of LTP in hippocampus**

The long-term potentiation of fEPSPs in the CA1 synapses was induced by standard protocol of high-frequency stimulation (HFS, 100Hz, 1 s-long) in the wild-type (A,B) and dnSNARE (C) mice kept in standard housing (SH, *left column*) or in an enriched environment (EE, *right column*). The average level for control WT mice is indicated as red line in all graphs.

(A) the potentiation of fEPSPs in the WT SH mice was induced during perfusion of individual astrocytes, lying in the vicinity of recording site, with intracellular solution containing either TexasRed dye alone (control) or TexasRed and TetanusToxin light chain or VGLUT inhibitor Rose-Bengal control (similar to Fig.5A). The inhibition of vesicular release of gliotransmitters from individual astrocytes significantly reduced the magnitude of LTP.

(B) Inhibitors of TREK-1 (spadin) and Best1 channels (CaCCInh-A01) had only marginal effect on the hippocampal LTP in the SH mice. The magnitude of LTP considerably increased in the EE mice. Inhibition of channel-mediated release of glutamate had more profound effect on the LTP magnitude than in the SH mice.

(C) Deficit in the astroglial exocytosis in the dnSNARE SH mice led to impairment of the LTP (similar to the data of Pascual et al. 2005). The application of exogenous D-Serine alone rescued only the short-term potentiation but not the LTP. However, the LTP phenotype was rescued by combining the D-Serine with a non-hydrolysable ATP analogue (both transmitters were applied for 5 min prior and 5 min after HFS). This result supports the importance of both the vesicular release of ATP and the non-vesicular release of D-Serine for regulation of NMDAR-dependent LTP.

The environmental enriched dnSNARE mice showed a much higher magnitude of LTP in area CA1, this results is consistent with our data on the LTP in the neocortex and results of the working memory test (Fig.6). The EE-induced enhancement of LTP was occluded by inhibition of either TREK-1 (spadin) or Best1 (CaCCInh-A01) channels.

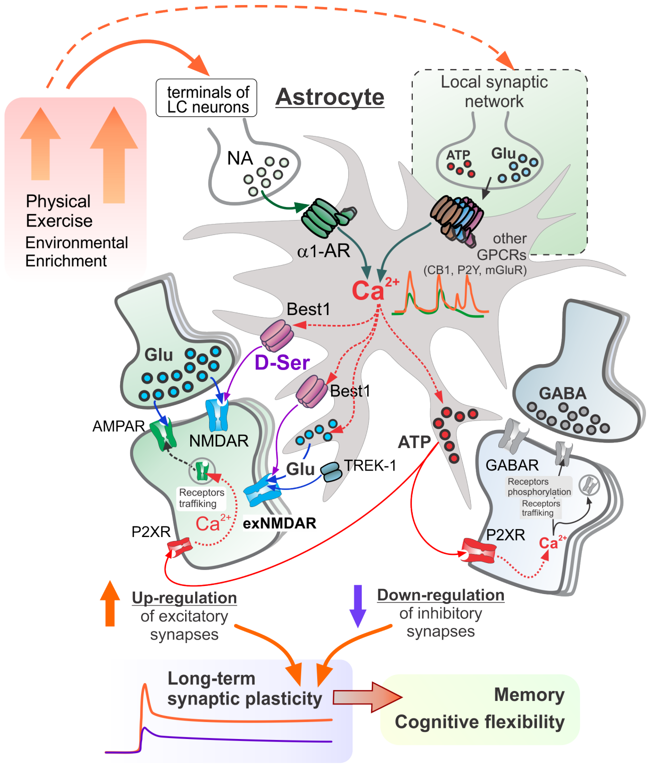


**Figure S6. Putative mechanisms of regulation of synaptic signalling and plasticity in neocortex by astrocytes**

Astrocytes release Astrocytes act a "Brain Hub" receiving signals from neurons and regulating synaptic transmission, neuronal metabolism and neurovascular coupling. Astroglial $Ca^{2+}$ signalling is activated by release of glutamate and other transmitters from local synapses and noradrenergic "volume transmission" from *Locus Coerules* axons (Bazargani and Attwell, 2016; Pankratov and Lalo, 2015; Savtchouk and Volterra, 2018).

<u>The synergy between vesicular and non-vesicular gliotransmitter release:</u>

- Elevation of $Ca^{2+}$ level in astroglial microdomains activates the Best1 channel-mediated release of D-Serine and exocytosis of glutamate and ATP (Lalo et al. 2014)
- Astroglial G-proteins also activate release of glutamate via TREK1 channels (Woo et al. 2012)
- Astroglia-derived D-Serine facilitates postsynaptic GluN2A-contaning NMDARs
- Best1-mediated release of D-Serine and vesicular release of glutamate from astrocytes activate extrasynaptic GluN2B-contaning NMDA receptors (exNMDARs), which manifests in the fast glia-induced inward currents (fGICs) in cortical neurons
- Best1-mediated release of D-Serine and TREK1-mediated release of glutamate activate slow glia-induced inward currents, mediated by exNMDARs
- Vesicular release of ATP from astrocytes activate extrasynaptic P2X receptors (Lalo et al. 2014) which up-regulate AMPAR-mediated synaptic signalling and down-regulate GABAergic inhibition via $Ca^{2+}$- and phosphorylation-dependent mechanisms (Boue-Grabot and Pankratov, 2017)
- Astroglia-driven up-regulation of glutamatergic and down-regulation of GABAergic signalling converge into the facilitation of long-term synaptic plasticity and working memory

Environmental enrichment and physical exercise enhance adrenergic signalling in astrocytes thereby increase $Ca^{2+}$-activated release of D-Serine, Glutamate and ATP which in turn facilitates experience-induced metaplasticity and cognitive flexibility.
